# Mapping RNA structure assembly and remodeling in biomolecular condensates

**DOI:** 10.64898/2026.05.21.726738

**Authors:** Ritwika Bose, Erik W. Bergstrom, Sovanny R. Taylor, Steven Boeynaems, Joshua A. Riback, Furqan M. Fazal, Anthony M. Mustoe

## Abstract

Biomolecular condensates concentrate RNA and protein machinery that facilitate RNA processing, ribonucleoprotein assembly, and gene regulation. Some evidence supports that condensates can modulate RNA folding, but measuring RNA structure within native condensates remains an unsolved challenge. Here we introduce RAID-MaP, a strategy that combines APEX proximity labeling with dimethyl sulfate (DMS) chemical probing to measure RNA structure within defined subcellular compartments. We applied RAID-MaP to resolve late stages of ribosomal RNA (rRNA) folding within the granular component (GC) of the nucleolus, revealing that both the 18S and 28S rRNAs feature widespread differences in structure compared to assembled ribosomes consistent with ongoing folding of both secondary and tertiary structure. We further combined RAID-MaP with transcription inhibition to resolve kinetics of rRNA maturation. The maturation kinetics of both subunits progressed on comparable timescales but with characteristic domain-level ordering, with late GC-resident intermediates often becoming more protected than mature ribosomes suggestive of stabilization by nucleolar accessory factors. Perturbing this 28S assembly pathway using antisense oligonucleotides produces distinct nucleolar phenotypes depending on whether early versus late folding domains are disrupted, demonstrating a direct link between rRNA folding and phase separation. We additionally applied RAID-MaP to discover that the 7SK small nuclear RNA undergoes spatially regulated structural switching consistent with localized release of the transcription factor P-TEFb at sites of active transcription. Together, our results establish subcellular spatial control of RNA structure as a new dimension of RNA regulation.

## Introduction

RNA molecules inhabit diverse cellular environments throughout their lifecycles, beginning with transcription and processing within specialized nuclear foci followed by localization of mature RNAs to varied cellular compartments^1,2^. Within each of these environments, differences in protein composition and physicochemical properties have the potential to modulate RNA folding^3,4^. However, the potential spatial variability of RNA structure within cells and its relationship to subcellular functional specialization remains poorly understood.

Emerging evidence points to deep relationships between biomolecular condensates and RNA folding. Condensates feature unique physicochemical environments, including elevated concentrations of RNA and RNA-binding proteins which may promote RNA structure remodeling relative to the surrounding milieu^5–9^. Reciprocally, RNA sequence and structural features can directly shape condensate organization^10–14^. A prominent example of this interplay occurs during ribosome biogenesis within the nucleolus, a nested condensate where nascent ribosomal RNA (rRNA) transcripts progressively fold and assemble with ribosomal proteins^15–19^. Current models of nucleolar organization posit that early pre-ribosomal intermediates engage in high-valency interactions with scaffolding proteins such as Nucleophosmin 1 (NPM1), and that as rRNA folds and matures, the progressive reduction in interaction valency allows export of assembled subunits^20,21^. However, evaluating this model is difficult due to our inability to directly measure rRNA folding within the native nucleolus. *Ex vivo* studies have reported high-resolution structures of many rRNA assembly intermediates^22–26^, but large fractions of the rRNA remain unresolved, and these discrete structures likely only reflect a subset of the branched assembly pathways that exist *in situ*. Thus, the precise relationship between rRNA folding and nucleolar form and function remains an open question.

Chemical probing experiments using reagents such as dimethyl sulfate (DMS) have widely enabled RNA structural characterization in living cells^27–29^. However, these experiments report population averages across the entire cell, masking potential structural changes that occur in minority states, such as RNAs partitioned into condensates. While genetic perturbations^30^, biochemical fractionation^31^, immunoprecipitation^32,33^, and metabolic labeling^34^ can be used to enrich specific RNA species for chemical probing or other structural analyses, these strategies typically disrupt native cellular organization, have limited generalizability, and lack spatial control. Here, we introduce RNA APEX Integrated with dimethyl sulfate (DMS) measured by Mutational Profiling (RAID-MaP) as a new method to probe RNA structure within native condensates. We apply RAID-MaP to resolve the assembly process of the human ribosomal RNA *in situ* in the nucleolus and structural remodeling of the 7SK small nuclear RNA across nuclear compartments. Our results reveal condensate-specific RNA structural remodeling as an important mechanism of RNA assembly and regulation and establish RAID-MaP as a broadly applicable platform for understanding how subcellular environments shape RNA structure.

### DMS and APEX chemistries are compatible and permit high-fidelity structure probing

RAID-MaP combines DMS chemical probing with APEX proximity labeling to enable measurement of RNA structure within targeted subcellular compartments (Fig. 1A). DMS globally modifies structurally accessible nucleotides throughout the cell^35^, while a genetically encoded APEX construct concurrently biotinylates RNA at a specific cellular location^1^. RNA is then harvested, biotin-enriched, and DMS modifications are quantified by mutational profiling (MaP) reverse transcription and deep sequencing. Structural changes unique to the APEX-targeted compartment are resolved by comparison to matched unenriched (bulk) DMS datasets. Importantly, DMS and APEX chemistries are orthogonal: DMS predominantly modifies the N1 and N3 positions of adenosine and cytidine, whereas the phenoxy-biotin radical generated by APEX primarily modifies the C8 position of guanosine (Fig. S1A)^1,36^. Both reactions are also rapid, permitting a combined 3-minute labeling time that minimizes RNA diffusion between compartments during the experiment.

**Figure 1.**
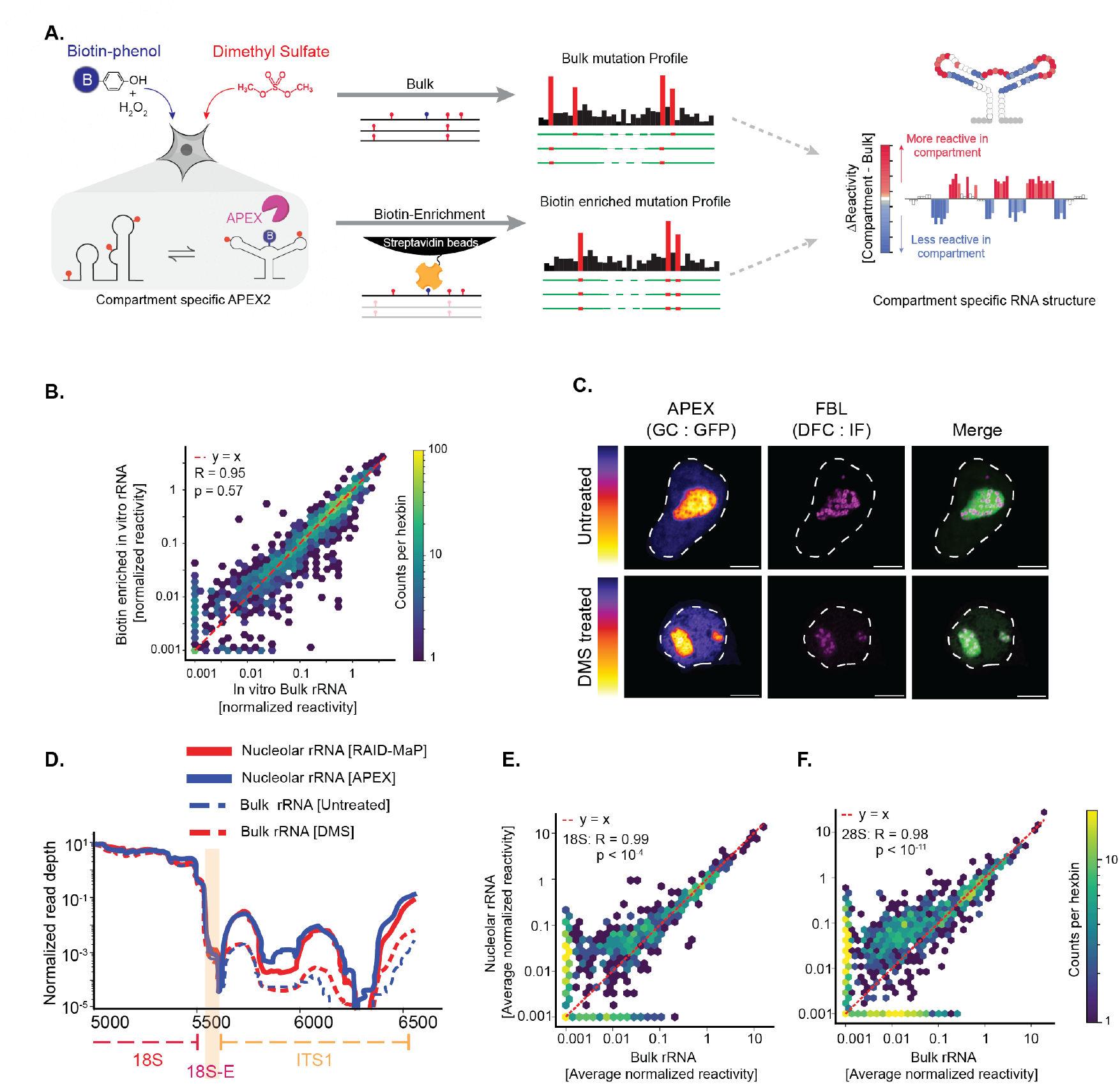
RAID-MaP enables subcellular RNA structure probing with nucleotide resolution. **A**. RAID-MaP workflow. Cells expressing APEX2 targeted to a defined subcellular compartment are treated with biotin-phenol (BP), H_2_O_2_, and dimethyl sulfate (DMS), enabling simultaneous proximity biotinylation and chemical probing. Extracted RNA undergoes mutational profiling (MaP) and high-throughput sequencing. Compartment-enriched ΔReactivity (ΔR) profiles are computed relative to bulk (unenriched) controls. **B**. Comparison of per-nucleotide DMS reactivities measured from E. coli rRNA treated with DMS, HRP, biotin-phenol, and H_2_O_2_ *in vitro* before and after biotin enrichment. Color scale, number of nucleotides per bin. P-value denotes the result of a two-sided Mann-Whitney U test comparing differences in magnitude of the two samples. **C**. Confocal immunofluorescence of cells expressing GC-targeted APEX2 (GFP, fire LUT) and the DFC marker fibrillarin (anti-FBL, magenta). Scale bar, 5 µm. **D**. GC enrichment captures non-transcribed regions of the 47S precursor. Normalized per-nucleotide read depth shows selective enrichment of ITS1 in GC-enriched relative to bulk samples, consistent with retention of pre-rRNA processing intermediates. The 18S-E sequence, which is retained until final processing in the cytoplasm, is highlighted. **E**., **F**. Hexbin plots comparing per-nucleotide DMS reactivity from bulk versus GC-enriched samples for 18S (E) and 28S (F) rRNA. Nucleolar reactivities are systematically elevated, indicating increased flexibility. Color scale, number of nucleotides per bin. P-value denotes the result of a two-sided Mann-Whitney U test comparing differences in magnitude of bulk and RAID-MaP measurements.

To validate the compatibility of DMS and APEX chemistries for RNA structural measurements, we performed *in vitro* probing experiments on model RNAs (extracted *E. Coli* rRNA and an *in vitro* transcribed adenosylcobalamin riboswitch) using DMS and the APEX analog horseradish peroxidase (HRP), which generates phenol radicals identical to APEX. Co-treatment with DMS and HRP had minimal impact on biotinylation compared with HRP alone (Fig. S1B). DMS-MaP libraries prepared from RNA that was HRP treated and biotin-enriched demonstrated modestly increased mutation rates in both DMS and ethanol controls compared to non-HRP-treated RNA (Fig. S1C), suggesting biotin modifications may impact RT fidelity. Standard normalization against HRP-alone samples effectively adjusted for these differences, yielding A and C reactivities that were highly correlated with those from DMS-only treated RNA (R > 0.95; Fig. 1B, S1D, S1E, S1G) and demonstrated equivalent accuracy for mapping secondary structure (AUROC = 0.89 and 0.74-0.78 for *E. coli* rRNA and AdoCbl riboswitch, respectively; Fig. S1F). These data confirm that high-fidelity RNA structure measurements can be obtained from RNA labeled with both APEX and DMS.

### RAID-MaP enables *in situ* probing of late ribosome assembly in the nucleolus

As a first application, we sought to use RAID-MaP to resolve the folding process of the human ribosome in the nucleolus. We focused on the stages of rRNA folding occurring in the outermost nucleolar layer, the granular compartment (GC), using an established APEX construct that concentrates in the GC within HEK293T cells^1^. Confocal microscopy confirmed APEX localization to the GC and that localization was minimally perturbed by DMS treatment (Fig. 1C). We obtained four replicates of matched bulk and biotin-enriched (henceforth termed nucleolar) RAID-MaP datasets for the 18S and 28S rRNAs. Nucleolar samples demonstrated reproducible ∼12-fold enrichment of reads mapping to external and internal transcribed spacer (ETS and ITS) precursor rRNA regions, confirming successful enrichment of immature rRNA species that only reside within the nucleolus (Fig. 1D, S2A-B)^37,38^. By contrast, the 18S-E precursor sequence, which is retained until final maturation in the cytoplasm^25,39^, was only enriched 1.7-fold (Fig. 1D, S2B). Junctions cleaved in the GC were more represented then those that occur in the dense fibrillar center (DFC)^19^, whereas later cleavage sites that mark progression toward the nucleoplasm and cytoplasm showed lower enrichment (Fig. S2C). Thus, nucleolar RAID-MaP enriches precursor rRNA intermediates relative to bulk measurements in a manner consistent with capture of GC-proximal stages of rRNA maturation.

RAID-MaP DMS reactivity profiles were highly reproducible across biological replicates (R>0.95; Fig. S2E) and provided coverage across ∼99% and ∼65% of the 18S and 28S rRNA respectively, with only a limited number of GC-rich expansion segments failing coverage cutoffs (Fig. S2F). Averaged nucleolar DMS reactivities were well-correlated with bulk reactivities but revealed distinctive signatures indicative of altered structural states within the nucleolus (Fig. 1E-F). Specifically, nucleotides within nucleolar 18S and 28S rRNAs displayed significantly higher average reactivities (P<0.001, Mann-Whitney U; Fig. 1E,F), consistent with increased conformational flexibility and less folded structure. Nucleolar reactivities also demonstrated modestly reduced agreement with mature rRNA secondary structure (AUROC = 0.71 vs 0.74; Fig. S2D). Note that despite a ∼12-fold nucleolar enrichment (Fig. S2B), the high abundance of mature rRNA means that 20–50% of ‘nucleolar’ samples still consist of mature ribosomes. For subsequent analyses, we thus focus on differential DMS reactivities, ΔR, representing the difference between APEX-enriched and bulk ribosomal subunits (Methods). Nonetheless, these data indicate that rRNA structure is, on average, destabilized in the GC and demonstrate that RAID-MaP can be applied in living cells.

### RAID-MaP recapitulates landmarks of small subunit assembly

We first examined structural differences observed in the 18S rRNA. Prior studies have suggested that the small subunit is substantially folded by the time it enters the GC^16,19^. Analysis of nucleotide-resolution ΔR values revealed distributed structural remodeling across the 18S rRNA (Fig. 2A). As expected, ΔR values are skewed towards positive ΔR values, indicating greater reactivity in the nucleolus that is consistent with reduced structure. However, significant numbers of negative ΔR nucleotides are also observed, indicating lesser reactivity that could reflect bound accessory factors or alternative local folding in the nucleolus. These ΔR differences predominantly map to nucleotides that are single-stranded in the assembled ribosome, consistent with ongoing maturation of 18S tertiary and quaternary structure (Fig. 2B). While paired nucleotides feature significantly fewer ΔR differences (Fig. 2B; P<10^-20^, Fligner-Killeen test), they still exhibit a significant skew towards positive ΔR (P<10^-25^; Wilcoxon signed-rank test). Thus, our data are consistent with the 18S rRNA featuring a largely folded but more dynamic secondary structure with significant tertiary structure heterogeneity in the GC.

**Figure 2.**
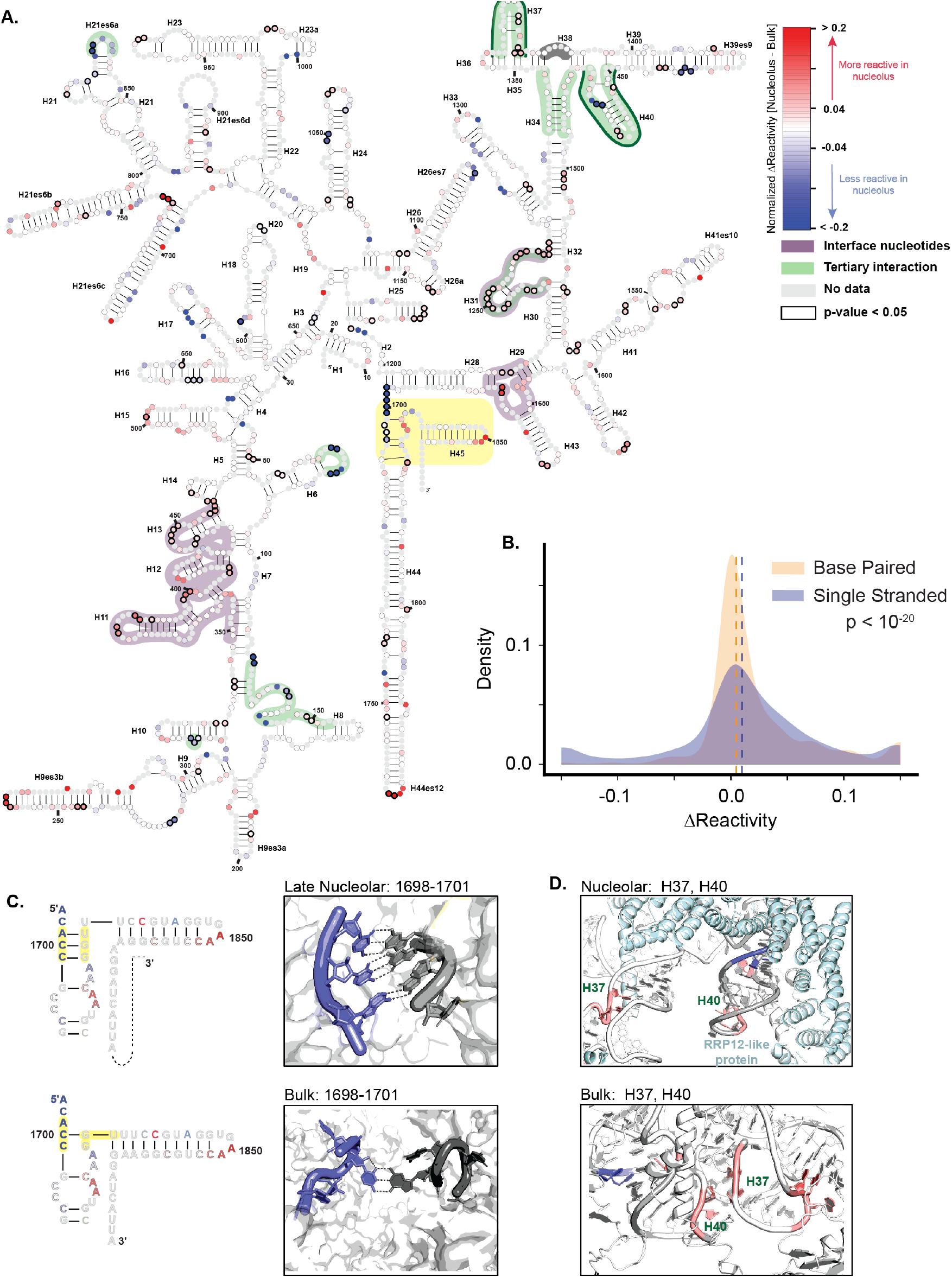
RAID-MaP resolves ongoing 18S rRNA assembly in the GC. **A**. Window-averaged ΔR across the 18S rRNA. Regions with increased (red) or decreased (blue) reactivity averaged over window of 3 nucleotides are plotted on a symmetric log (symlog) scale. Black outlined circles denote nucleotides with statistically significant differences (p < 0.05, Wilcoxon-rank sum test). Nucleotides involved in 18S-28S subunit interfaces are highlighted in purple, and those participating in long-range tertiary contacts are shown in green. The H37–H40 tertiary interaction (dark green border) is expanded in panel D. Helices H44 and H45 (yellow shading) are expanded in panel C. **B**. ΔR is enriched at single-stranded nucleotides. Kernel density estimate (KDE) plots compare ΔR distributions at paired versus unpaired sites in the mature ribosome, revealing greater variance at unpaired positions. Significance was assessed using a Fligner-Killeen test. Matched colored dashed lines indicate the median of each distribution. **C**. Detail of the H44/H45 region showing secondary structure in immature and mature ribosomes with ΔR values superimposed. Cryo-EM structures of the mature ribosome (PDB 6QZP) and late nucleolar pre-40S particle (PDB 6G18) are shown for comparison. **D**. Structural comparison of H37 and H40 in mature 80S (PDB 6QZP) and nucleolar pre-40S (PDB 7MQ8) ribosomes. Nucleotides with significant ΔR are highlighted; neighboring ribosomal proteins are shown in white. RRP12-like protein (cyan), present only in the pre-40S particle, contacts H37 and H40 and maintains them in an immature conformation.

To better pinpoint major GC-specific structural differences, we used a rolling 3-nt window to identify regions with significant local structural changes (black highlight, Fig. 2A,C). 131 regions exhibit significantly positive ΔR, and 96 regions with negative ΔR (Fig. 2A). Many of these changes recapitulate known features of immature small subunits. For example, regions that form contacts with the large subunit displayed significantly increased ΔR, including loops in H11–15, H28-29, and H31 (Fig. 2A, purple shading). The apical loop of H44 also showed elevated reactivity in the nucleolus, consistent with H44 being undocked^23,40,41^. By contrast, the base of H44 (nts 1697–1705) exhibited reduced reactivity, consistent with alternative base-pairing previously observed in nucleolar intermedates^39,40,42^ (Fig. 2C). Multiple long-range tertiary contacts in the mature 18S also exhibit significantly increased ΔR, denoting immature folding. For example, we observe significant ΔR in H37 and H40, consistent with high-resolution structures of state E where H37 and H40 are held open by RRP12-like protein^26,43^ (Fig. 2D). These data thus provide direct evidence of correspondence between isolated assembly intermediates observed by cryo-EM and native assembly in the nucleolus.

### 28S rRNA features greater structural heterogeneity in the GC

Compared to the 18S rRNA, less is known about 28S folding pathways, with only a minority of 28S rRNA nucleotides resolved in structures of large subunit assembly intermediates (Fig. S2F). We performed an identical ΔR analysis across the 28S sequence (Fig. 3A, B). As with the 18S, ΔR differences predominantly map to nucleotides that are single-stranded in the mature large subunit (Fig. 3B; P<10^-29^, Fligner-Killeen test). However, base-paired nucleotides also show significantly elevated reactivities in the 28S compared to 18S (P<10^-8^, Mann-Whitney U), indicating greater heterogeneity of 28S rRNA secondary structure in the nucleolus. Notably, rolling 3-nt window analysis revealed that 41% of statistically significant ΔR regions mapped to nucleotides that are base-paired in the mature 28S rRNA, with the majority of these (80%) exhibiting increased nucleolar reactivity. Representative examples include nucleotides 2491-2499 in ES19, and nucleotides 1461-1463 within H30, indicating that these helices remain destabilized throughout GC steps of 28S rRNA assembly (Fig. 3A, S3). Together, these data indicate that a larger fraction of the 28S rRNA remains conformationally immature within the GC relative to the 18S, encompassing both secondary and tertiary structural elements.

**Figure 3.**
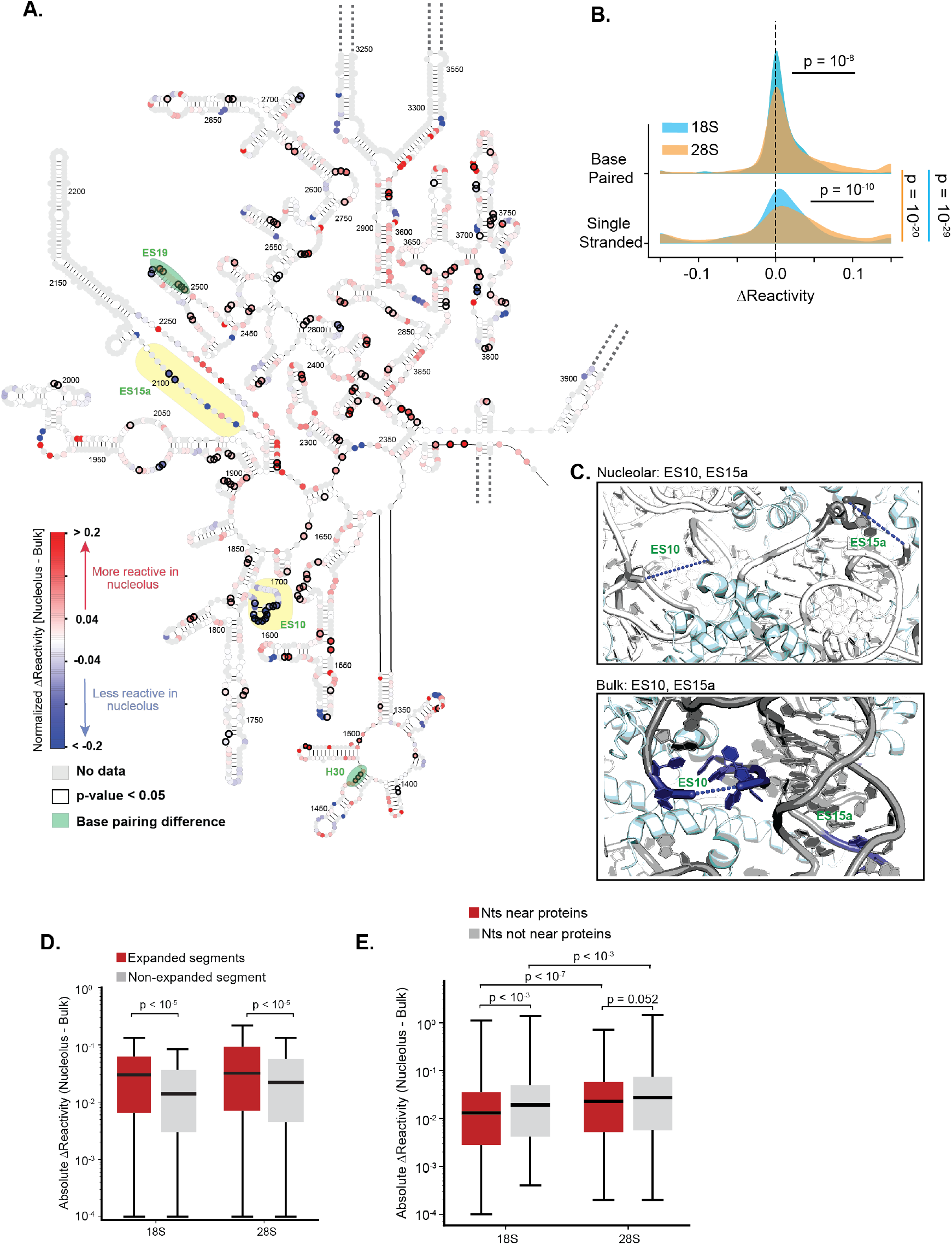
28S rRNA exhibits greater structural heterogeneity than 18S rRNA in the GC. **A**. Windowed mean ΔR for domains II, III, and IV of the 28S rRNA plotted on a symlog scale. Yellow highlights a proposed tertiary interaction between ES10 and ES15a (expanded in panel C); green highlights nucleotides with elevated reactivity at nominally base-paired positions. **B**. ΔR is enriched at single-stranded nucleotides in 28S rRNA. KDE plots comparing ΔR at paired versus unpaired sites, showing greater dispersion at unpaired positions. P values for differences in dispersion between paired and unpaired nucleotides within each rRNA were calculated using the Fligner–Killeen test. Differences in the distributions of ΔR values between paired and unpaired nucleotides were assessed using the Mann–Whitney U test. **C**. Cryo-EM structures of mature (PDB 6QZP) and late nucleolar (PDB 8FKX) 28S rRNA colored by RAID-MaP ΔR. Dashed lines indicate segments unresolved in cryo-EM. Ribosomal proteins shown in cyan. **D**. Expansion segments undergo greater structural remodeling than core regions. Box plots of ΔR at expansion segment versus core nucleotides in both subunits. Significance calculated using Mann–Whitney U test. **E**. 28S rRNA sites near ribosomal proteins are more flexible than their 18S counterparts. Box plots of ΔR at nucleotides within 5Å of ribosomal protein contacts in 18S and 28S. Significance calculated using Mann–Whitney U test.

We observed multiple regions with pronounced decreases in nucleolar reactivity, suggestive of formation of alternative structural states or protection by nucleolar-specific assembly factors (Fig. S3). The most prominent example is ES10, where ten consecutive nucleotides show significantly reduced reactivity in the nucleolus (Fig. 3C). A cluster of strongly decreased reactivities were also observed in ES15, which is located in close proximity to ES10 (Fig. 3C). Notably, ES10 was shown to be essential for late subunit assembly in yeast, but via an unknown mechanism^44^. The strong protections observed via RAID-MaP suggest that this region may function as a binding site of nucleolar-specific assembly factors. More broadly, nucleotides in expansion segments (ESs) exhibit significantly more reactivity changes in the GC than non-ES nucleotides (Fig. 3D; P<10^-5^, Mann–Whitney U). This is consistent with ESs structures being among the last to consolidate during large subunit assembly, possibly due to transient engagement by nucleolar assembly factors that prevent stable ES folding.

We also examined whether the reduced stability of 28S rRNA was related to delayed incorporation of ribosomal proteins (r-proteins). In the 18S rRNA, nucleotides located within 5 Å of ribosomal proteins in the assembled ribosome showed reactivities significantly closer to bulk then those at more distal nucleotides (Fig. 3E; P<10^-3^, Mann–Whitney U), suggesting that many r-proteins are incorporated into the small subunit prior to its entry to the GC. By contrast, protein-proximal and protein-distal nucleotides demonstrated statistically insignificant ΔR distributions in 28S rRNA, suggesting that the GC is a major site of r-protein incorporation into the 28S (Fig. 3E). Similar analysis of ΔR based on proximity to post-transcriptional modification sites revealed no relationship, consistent with most modifications being installed prior to 18S and 28S trafficking to the GC^19^ (not shown).

Together, these results indicate that the 28S rRNA features pronounced structural differences in the GC including at the secondary, tertiary, and quaternary structural level, consistent with the GC playing a key role in large subunit folding and r-protein incorporation.

### rRNA domains exhibit heterogeneous maturation kinetics

Studies in bacteria, yeast, and human cells have indicated that rRNA folding and assembly is loosely ordered by domains^23,45,46^. Analysis of ΔR by domains failed to reveal a clear ordering, with all domains exhibiting broad ΔR distributions (Fig. 4A, C). However, this is unsurprising given that steady-state RAID-MaP yields an average over many assembly states that exist in the GC. A leading model of the GC is that rRNA assembly and phase separation are intimately coupled: early intermediates flowing in from the DFC form high valency interactions with GC scaffolding proteins such as NPM1, and the valency of these rRNA-scaffold interactions then decreases as the rRNA progressively folds, resulting in eventual release of mature subunits into the nucleoplasm^20^. Thus, the steady-state GC population likely reflects an enrichment of early, high-valency intermediates. To test this model and better resolve potential assembly hierarchies, we pursued a temporal perturbation strategy where we inhibited new rRNA transcription with low concentrations of actinomycin D (ActD), effectively reducing the influx of new intermediates from the DFC without impacting maturation of GC-resident intermediates^47,48^. Notably, because only molecules that persist in the GC are captured by RAID-MaP, successive timepoints provide kinetic resolution on domain-specific maturation along with insights into nucleotide-level features that underly rRNA phase separation in the GC.

**Figure 4.**
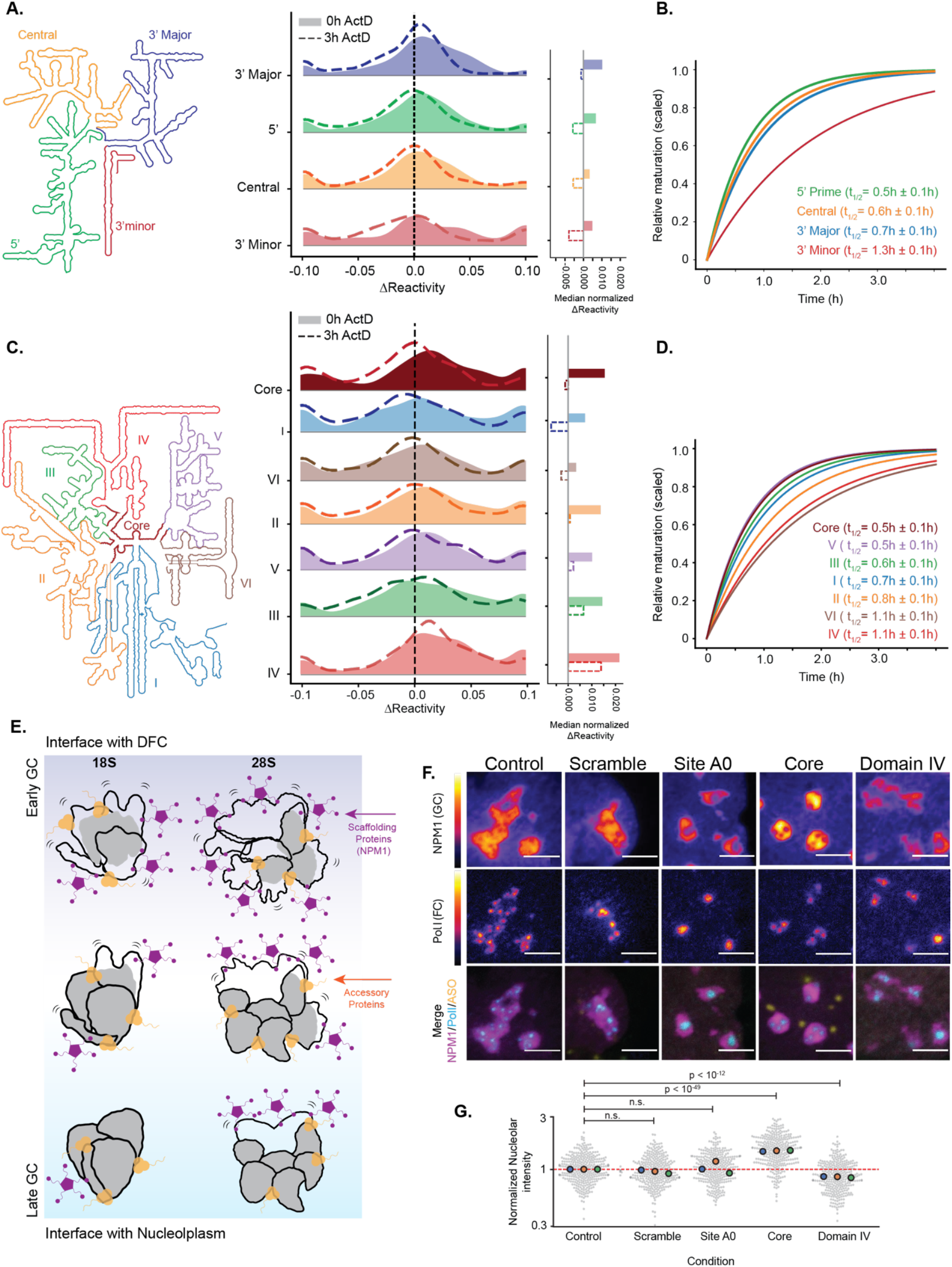
Domain-ordered rRNA maturation within the GC governs nucleolar condensate architecture. **A**. Domain-level ΔR for 18S rRNA during ActD treatment. Schematic of the four canonical 18S domains (5′, Central, 3′ Major, 3′ Minor; color-coded). Right, ΔR distributions at 0 h (solid) and 3 h actinomycin D (ActD; dashed). Vertical dashed line, ΔR = 0. Bar plots, median normalized ΔR per domain. **B**. 18S domains mature with distinct kinetics. Cumulative normalized ΔR over the ActD time course for each domain in A. Exponential fits yield domain-specific half-lives (t1/2). **C**. Domain-level ΔR for 28S rRNA during ActD treatment. Schematic of the six structural domains (I–VI) and central Core of 28S rRNA. Right, ΔR distributions at 0 h (solid) and 3 h ActD (dashed). Vertical dashed line, ΔR = 0. Bar plots, median normalized ΔR per domain. **D**. 28S domains mature with distinct kinetics. Cumulative normalized ΔR over the ActD time course for each domain in C. **E**. Model of rRNA structural maturation and GC retention for the 18S and 28S subunit precursors. The 18S rRNA (left) reaches near-mature secondary and tertiary structure within the GC, with retention mediated primarily by accessory factor (orange) occupancy at late stages. The 28S rRNA (right) retains substantial structural immaturity throughout GC residence, with incompletely folded domains sustaining prolonged scaffold protein (NPM1, purple) interactions. Color (gray) reflects the degree of structural completion relative to the assembled ribosome. **F**. ASO perturbation of rRNA folding induces distinct nucleolar morphologies. U2OS cells expressing endogenously tagged NPM1 (GC marker, fire LUT) and RNA Pol I (FC marker, fire LUT). Fluorescently labeled ASOs targeting the 28S core or domain IV are shown in yellow. Scale bar, 4 µm. **G**. Quantification of ASO-induced changes in GC NPM1 concentration. Swarm plots of NPM1 fluorescence intensity across ASO conditions, normalized to control. Each point represents an individual cell. Three independent replicates were performed, with per-replicate means plotted with color circles. Significance calculated using Welch’s t-test aggregating cells from all replicates together

ActD treatment induced characteristic nucleolar reorganization with rapid displacement of the FC and DFC while largely preserving the GC^15,49^ (Fig. S4A). We performed RAID-MaP experiments at 1, 2, and 3 hrs post ActD treatment. Nucleolar DMS reactivities demonstrated a general reduction in ΔR magnitude following a roughly exponential time course, consistent with a global maturation process (Fig. 4A,C, S4B). Because four timepoints was generally insufficient for robust nucleotide-level kinetic fitting, we quantified these dynamics at a coarse-grained, domain-level resolution, fitting grouped ΔR trajectories to an exponential model to yield per-domain t_1/2_ values. These values summarize the kinetics of how each domain shifts toward the final GC-enriched intermediate state after new rRNA synthesis is blocked (Methods).

For the 18S rRNA, the 5′, central, and 3′ major domains all demonstrated roughly equivalent kinetics of maturation following transcriptional arrest with ActD, with t_1/2_ = 0.5-0.7 ± 0.1h (Fig. 4B). These domains also demonstrate a substantial reduction in nucleotides with ΔR>0 and modest enrichment in nucleotides with ΔR<0 (Fig. 4A), indicating enhanced protection compared mature ribosomes that is suggestive of interactions with GC proteins. By comparison, the 3′ minor domain matured more slowly (t_1/2_ = 1.3h ± 0.1; Fig. 4B), and retained a much broader ΔR distribution and a pronounced negative tail (P<0.001 for all pairwise comparisons against the 3′ minor domain, Fligner-Killeen). Thus, these data indicate that assembly of the 3′ minor domain is decoupled from other 18S rRNA domains and is potentially stabilized in an undocked state by interactions with GC proteins.

For 28S rRNA, the central core and domains I, III, and V mature quickly at rates similar to the 18S, with t_1/2_ = 0.5-0.7 ± 0.1h (Fig. 4D). Notably, the fast maturation of the core and domain V are consistent with early establishment of the peptidyl transferase center observed in human pre-60S intermediates^24^. Domain II demonstrated intermediate kinetics (t_1/2_ = 0.8h ± 0.1), and domains IV and VI matured the slowest (t_1/2_ = 1.1h ± 0.2). Similar to the 18S, most domains demonstrate a reduction in nucleotides with ΔR>0 and modest enrichment in nucleotides with ΔR<0 at t=3h endpoint (Fig. 4C). However, domain III and particularly domain IV are exceptions, remaining substantially less structured than the mature ribosome at t=3h (ΔR>0; Fig. 4C), suggesting that these domains remain in a more heterogeneous, unfolded state.

Together, our ActD data indicate that the maturation of the 18S and 28S in the GC follow comparable kinetics, with most domains maturing on the half-hour timescale and a minority taking around an hour. Across both subunit intermediates, many nucleotides switch from higher to lower reactivity relative to the mature ribosome, consistent with a transition from more promiscuous interactions, such as those involving NPM1, toward more specific interactions with ribosome biogenesis factors^21^. For the 28S, however, this trend is domain dependent, consistent with previous reports that NPM1 enrichment differs between synthetic nucleoli formed with the 18S or 28S alone^19^. Altogether, these observations support an updated model for rRNA maturation within the GC (Fig. 4E).

### rRNA folding modulates nucleolar condensate architecture

We sought to orthogonally test the assembly hierarchy resolved by RAID-MaP using antisense oligonucleotides (ASOs) to perturb rRNA folding. Because the nucleolus is organized by multivalent interactions between pre-ribosomal particles, assembly factors, and scaffolding proteins^20,21,50^, we reasoned that we could use nucleolar phase separation as a proxy of rRNA folding status. Specifically, the concentration of the GC scaffolding protein NPM1 is expected to depend on the concentration of rRNA surfaces available for forming multivalent interactions. ASOs that trap early rRNA assembly should promote high rRNA valency, and therefore high NPM1 concentration, whereas trapping late assembly intermediates should result in low valency and low NPM1.

We transfected a series of steric blocking ASOs into U2OS cells expressing endogenously tagged NPM1-mCherry and Pol I-muGFP^51^, allowing visualization of FCs and quantification of relative NPM1 GC concentrations (Fig. 4F). ASOs contained a 5’ terminal alkyne allowing attachment of a fluorophore after cell fixation to confirm ASO uptake and localization (Fig. 4F). As a positive control, we first used a previously published^52^ ASO that blocks the pre-rRNA cleavage site A0, an early processing step required for downstream 18S rRNA maturation. As expected, this A0 ASO (ASO_A0_) induced enlarged FC/DFC structures, with a minor decrease in GC size but no impact on NPM1 concentration in the GC (Fig. 4F, G). We next used ASOs to disrupt the 28S core domain (ASO_core_) and domain IV (ASO_IV_), which RAID-MaP indicated were early and late assembly steps, respectively. Strikingly, these two ASOs had minimal impact on FC/DFC size but caused dramatically different GC phenotypes. ASO_core_ caused rounding of the GC and increase in NPM1 concentration (p < 0.01, Welch’s t-test, Fig. 4F, G), consistent with a block of early assembly that promotes accumulation of immature, high-NPM1-valency 28S intermediates. By contrast, ASO_IV_ induced fragmentation of the GC and reduction in NPM1 concentration (p < 0.01, Fig. 4F, G), consistent with accumulation of more mature, low-NPM1-valency 28S intermediates.

Together, these findings support the 28S assembly hierarchy resolved by RAID-MaP and provide direct evidence that rRNA folding and valency underpin GC phase separation^20^.

### 7SK RNA undergoes localized structural remodeling at sites of active transcription

We finally sought to extend RAID-MaP to investigate location-specific structural remodeling of other RNAs. We focused on the human 7SK non-coding RNA, which plays a central role in regulating transcription elongation by chaperoning the transcription factor P-TEFb (positive transcription elongation factor b; also known as CDK9/Cyclin T)^53^. Bulk chemical probing studies have shown that 7SK encodes a large-scale structural switch, populating at least three distinct structures in cells that correspond to P-TEFb bound, released, and potential intermediate states, respectively^54,55^. 7SK exists broadly in the nucleoplasm, but also partitions into nuclear speckles, which are condensates that coordinate transcription and splicing^56^ (Fig. S5A). Specifically, evidence indicates that P-TEFb is released from 7SK at nuclear speckles to locally activate transcription at speckle-proximal genes^57,58^, but the relationship between 7SK structure, P-TEFb release, and spatial control of transcription remains unknown.

We used RAID-MaP to probe 7SK structure in nuclear speckles using an established SRSF1– APEX2 fusion expressed in HEK293T cells^59^. As a comparison, we also probed local 7SK structure in the nuclear lamina, a transcriptionally-repressed nuclear compartment, using a LaminA/C-APEX2 fusion construct^1^. Confocal microscopy confirmed intended localization of each APEX-fusion (Fig. 5A), and qPCR following DMS treatment, APEX labeling, and biotin enrichment confirmed successful enrichment of nuclear transcripts, whereas cytoplasmic transcripts were depleted (Fig. 5B). Amplicon sequencing of 7SK yielded bulk DMS profiles consistent with previously reported 7SK datasets^60^ (R = 0.81; Fig. S5B), with minor differences within the range expected from experimental variation in DMS probing (Methods). DMS reactivity profiles for both enriched and matched bulk samples were highly reproducible across 2-3 biological replicates (R>0.95; Fig. S5C). Importantly, single-molecule DANCE deconvolution of bulk DMS data recovered the previously defined 7SK states A, B, and H, corresponding to P-TEFb bound, released, and putative intermediates states, respectively (Fig. S5D)^54^. Single-molecule deconvolution was infeasible on biotin-enriched samples due to low yields of unique 7SK reads after sequencing and deduplication; we therefore used a pseudoinverse decomposition strategy to estimate 7SK state populations by fitting against the A, B, and H state reactivities determined from the deconvolved bulk sample (Fig. 5D). Despite only modest apparent differences in the averaged reactivity profiles of speckle- and lamina-enriched 7SK (Fig. 5C), this analysis revealed a pronounced redistribution of 7SK conformational states across compartments. Speckle-enriched samples demonstrate strong enrichment of State B relative bulk (0.53 vs. 0.35), consistent with localized release of P-TEFb. By contrast, lamina-enriched samples were strongly enriched for State A (0.58 vs. 0.43), consistent with P-TEFb being bound and inhibited in the transcriptionally repressed lamina environment.

**Figure 5.**
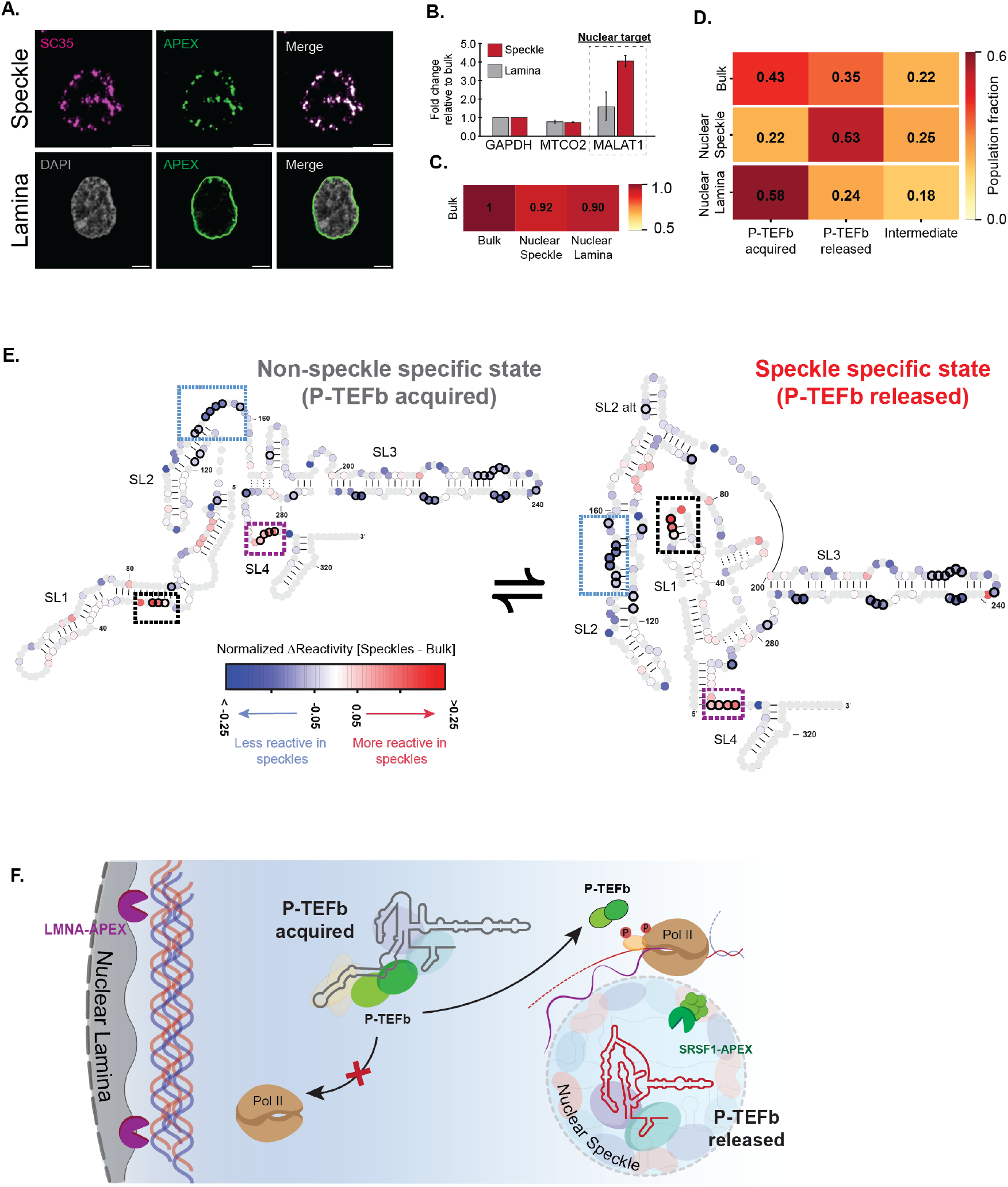
7SK structure is spatially regulated in the nucleus. **A**. Confocal immunofluorescence of cells expressing SRSF1-APEX2 (anti-FLAG, green) co-stained with the speckle marker SC35 (magenta), or LMNA-APEX2 co-stained with DAPI. Scale bar, 4 µm. **B**. qPCR quantification showing enrichment of the nuclear-retained RNA MALAT1 over cytoplasmic transcripts in DMS treated, biotin-enriched fractions from SRSF1-APEX2 and LMNA-APEX2 cells. **C**. Pearson correlation heatmap comparing mean normalized DMS reactivities between bulk, speckle-enriched, and lamina-enriched 7SK. Means were computed from n = 3, n=3, and n=2 experiments for bulk, speckle, and lamina samples, respectively. **D**. 7SK state populations in each nuclear compartment estimated by pseudoinverse decomposition against DANCE–deconvolved bulk state reactivities. **E**. 7SK RNA adopts distinct structural ensembles in nuclear speckles relative to bulk. Windowed mean ΔR (enriched − bulk) from nuclear speckle RAID-MaP overlaid on the secondary structures of State A (left) and State B (right). Nucleotides more reactive in speckles (red) or less reactive in speckles (blue) are shown on a symlog scale (color bar). Black outlined circles denote nucleotides with statistically significant differences (p < 0.05, Wilcoxon-rank sum test). Dashed boxes highlight structural elements showing the most pronounced (black, magenta) or least pronounced (blue) compartment-specific differences. **F**. Model of compartment-specific 7SK conformational switching. State A (P-TEFb-bound and inhibited) predominates at transcriptionally repressed nuclear lamina. State B (P-TEFb-released) predominates at transcriptionally active nuclear speckles.

Detailed ΔR analysis corroborated these compartment-specific 7SK population shifts. In speckles, significantly increased ΔR is observed at nts 25–29, precisely localizing to the apical loop of SL1alt, a structure that is uniquely present in State B (Fig. 5E, S5E-F)^54^. Furthermore, nts 105– 120 and 140–160 exhibit significantly decreased reactivity, consistent with formation of SL2ext in State B. Increased reactivity is also observed at nts 295–299, which is a highly reactive loop in both States A and B. In contrast, lamina-enriched samples showed reductions in reactivity at nts 25–29, consistent with State A enrichment (Fig. S5E-F). Interestingly, the SL3 domain features significantly reduced reactivity in both speckles and lamina, indicating relative stabilization of SL3 (Fig. 5D, S5E-F). Because unfolding of SL3 is the defining feature of state H^54,60^, these observations indicate that the state H is likely depleted from speckles and lamina and thus implying that state H is associated with some other nuclear compartment. Together, these data demonstrate that 7SK populates distinct conformational states in different nuclear compartments that correlate with spatial organization of transcription.

## Discussion

Whether subcellular localization shapes RNA folding has been difficult to directly measure. We introduced RAID-MaP as a new strategy that combines APEX proximity labeling with DMS chemical probing to enable measurement of RNA structure with subcellular specificity at nucleotide resolution. Using RAID-MaP to probe human rRNA and 7SK across multiple locations reveals that both RNAs feature significant local regulation of RNA structure that is correlated with compartment-specific RNA biochemical function. Further, our studies of the nucleolar GC establish that RNA folding state can also reciprocally influence condensate architecture. These findings establish subcellular localization as a novel regulatory dimension of RNA structure, with important implications for our understanding of the form and function of biomolecular condensates.

Within the nucleolus, we resolve the domain-level maturation program of rRNA in the GC, the final compartment before subunit export. The ordering we uncover, with architectural core elements consolidating first and peripheral domains last, is broadly consistent with cryo-EM-derived assembly intermediates^24,40^. Consistent with previous reports^19,21,52^, our measurements indicate that 18S intermediates are more structurally mature than 28S intermediates in the GC. However, both subunits exhibit comparable global maturation kinetics within the GC upon ActD inhibition of transcription. Slower maturation kinetics of peripheral 18S 3' minor and 28S domains IV, as well as a general shift of most domains towards reduced reactivity compared to mature ribosomes, is consistent with formation of GC-specific protein interactions. Together with evidence that accessory factors to ribosome biogenesis contain conserved intrinsically disordered regions^21^, exhibit higher GC partitioning, and are associated with reduced local mesh size^50^, these observations support a model in which rRNA structural maturation remodels the interactions that retain pre-ribosomal intermediates in the GC. Early, structurally dynamic intermediates are retained through non-specific scaffold interactions with GC proteins, including NPM1. As rRNA folds, these scaffold interactions decrease, while IDR-containing accessory factors maintain multivalent GC retention until intermediates are sufficiently mature for release. Thus, evolving rRNA secondary and tertiary structure tunes the balance between non-specific scaffold- and accessory-factor-mediated retention within the phase-separated GC (Fig. 4E).

Our studies of 7SK reveal that the structural switch between P-TEFb bound and released conformations is spatially regulated in the nucleus (Fig. 5F). Nuclear speckles are generally located proximal to highly transcribed genes, whose chromatin loops extend to the speckle periphery^58^. The enrichment of the P-TEFb-released conformation of 7SK (State B) in speckles is therefore consistent with a population of 7SK that has completed P-TEFb handoff at the speckle periphery and is potentially poised for reloading. By contrast, 7SK molecules proximal the nuclear lamina predominantly exist in State A, suppressing P-TEFb activity at the transcriptionally suppressed lamina. It is important to note that our data are unable distinguish whether 7SK structural remodeling is directly promoted by speckle localization, or rather if P-TEFb-released 7SK preferentially partitions into speckles. Nevertheless, we posit that 7SK functions as a vehicle to deliver P-TEFb to targeted chromatin loci while preventing indiscriminate P-TEFb activity at non-target loci.

RAID-MaP has several important limitations. Spatial resolution is determined by the diffusion radius of RNAs over the course of the 2 min DMS probing time, and diffusion of APEX-generated biotin-phenoxyl radicals during the 1 min APEX labeling time, and thus likely also captures RNAs near compartment boundaries. Isolation of RNA after labeling is also imperfect, with substantial residual contamination of bulk RNA that contributes to the DMS reactivities measured by RAID-MaP. DMS data are also averaged over what may be many heterogeneous states within a given compartment. Finally, DMS only reports on whether nucleotides are accessible or protected and cannot distinguish the mechanism of protection (e.g. base pairing versus protein binding).

Overall, RAID-MaP represents a broadly applicable strategy for resolving mechanisms of RNA assembly and differential regulation that are compartmentalized with the cell. We envision expanding RAID-MaP to resolve RNA structural landscapes in other RNA-containing condensates and membrane-bound locales, and integrating with RAID-MaP with proximity proteomics to further link RNA structural remodeling to local protein environments. Our results establish localized remodeling of RNA structure as a likely key dimension of RNA function.

## Acknowledgements

We thank B. Blencowe (University of Toronto) for providing the SRSF1-APEX cell line. We also thank C. Rosenblad (Baylor College of Medicine [BCM]) for assistance with imaging, I. Saleem (BCM) and S. Sharma (BCM) for technical support, and S. Pradhan (BCM) for generating the LMNA-APEX cell line. A.M.M., F.M.F., J.A.R., and S.B. are CPRIT Scholars in Cancer Research. This work was supported by the Cancer Research and Prevention Institute of Texas (RR190054 to A.M.M.; RR210012 to F.M.F; RR210040 to J.A.R.; RR220094 to S.B.), National Institutes of Health (R35GM147010 to A.M.M.; R00HG010910 and R35GM154922 to F.M.F.; R35GM162528 to J.A.R.; DP2NS142714 and R01NS138605 to S.B.), the Welch Foundation (Q-2186-20240404 to F.M.F; Q-2254-20250403 to J.A.R.; Q-2180-20240404 to S.B.), the Searle Scholars Program (J.A.R.), NSF (DBI # 2213983, WALII to S.B.) and BCM seed funds (to F.M.F).

## METHODS

### RAID-MaP probing of adenosylcobalamin riboswitch

DNA template encoding AdoCbl was purchased as a gBlock (IDT), PCR-amplified using Q5 DNA polymerase (NEB), and purified with PureLink PCR columns (Invitrogen). *In vitro* transcription was performed with 250 ng of PCR product using HiScribe T7 High Yield RNA Synthesis kit (NEB), followed by DNase treatment (TURBO DNase, Invitrogen) and purification (Mag-Bind TotalPure NGS, Omega Biotek). RNA size and purity were confirmed by TapeStation analysis (Agilent).

For structure probing experiments, 2.4 µM RNA was denatured at 95°C for 2 min, placed on ice for 2 min, and refolded at 37°C for 30 min in folding buffer (1× buffer: 325 mM bicine (pH 8.3), 200 mM NaCl, 15 mM MgCl_2_, and 0.8 mM adenosylcobalamin (Sigma)), followed by addition of 2.7 µM Horseradish Peroxidase (ThermoFisher) and 0.5 mM biotin-phenol (Iris Biotech). DMS+biotin and ethanol+biotin reactions were performed by adding 0.16 M DMS in ethanol or 100% ethanol, incubating at 37 °C for 5 min, followed by addition of 0.1 mM H_2_O_2_ and incubation for 1 minute. For HRP-only conditions, 0.1 mM H_2_O_2_ was added directly and incubated for 1 minute. Reactions were quenched with an equal volume of 20% β-mercaptoethanol (BME) in H_2_O and then placed on ice. For each condition, control reactions excluding H_2_O_2_ were also performed to assess biotinylation specificity. Samples were purified using columns (Zymo RNA Clean & Concentrator) and eluted in 5 µL. For each condition, six reactions were performed in parallel and then pooled for down-stream analysis. Two independent experiments were performed on different days.

### RAID-MaP probing of cell-free *E. coli* rRNA

Protein-free total RNA from *E. coli* K-12 MG1655 was prepared as previously described^29^. Overnight cultures were grown in LB at 37°C to OD_600_ ≈ 0.5, pelleted, and resuspended in lysis buffer (15 mM Tris–HCl (pH 8), 450 mM sucrose, 8 mM EDTA (pH 8), 0.04 mg/mL lysozyme). RNA was extracted using three rounds of phenol/chloroform/isoamyl alcohol (PCA) followed by three rounds of chloroform extraction and eluted into folding buffer (1×: 200 mM Bicine (pH 8.3), 200 mM potassium acetate (pH 8.0), 0 or 5 mM MgCl_2_). After equilibration at 37°C for 10 min, 41 µg total RNA in 157 µL was combined 2.5 µM HRP and 0.5 mM biotin-phenol. Samples were treated with 0.16 DMS in ethanol or 100% ethanol and incubated at 37 °C for 5 min. Biotinylation was initiated by adding 0.1 mM H_2_O_2_ for 1 min, followed by quenching with 200 µL of ice-cold 20% BME. For each condition, control reactions excluding H_2_O_2_ were also performed to assess biotinylation specificity. RNA was purified (RNeasy Midi, Qiagen), DNase-treated (TURBO DNase, ThermoFisher), and purified again (RNeasy Midi, Qiagen) before downstream analysis.

### Biotin enrichment *in vitro* probed RNAs

Biotinylated adenosylcobalamin and cell-free *E. coli* RNAs were enriched by adapting a previously published protocol^61^. 30 µL of Dynabeads MyOne Streptavidin C1 (Invitrogen) beads were used per 10 µg of input RNA. Beads were washed three times in 300 µL of hybridization buffer (33.3 mM Tris-HCl (pH 7.0), 500 mM NaCl, 0.7 mM EDTA, 0.7 mM SDS, 10% formamide), incubating for 5 min at room temperature per wash. Beads were resuspended in 300 µL of hybridization buffer, mixed with input RNA (∼10 µL volume), and incubated at 37 °C for 30 min on a rotator. Beads were captured on a magnetic stand and the supernatant was removed. Beads were washed twice in 300 µL of wash buffer (10 mM Tris-HCl (pH 7.5), 1 mM EDTA, 2 M NaCl), incubating for 5 min at room temperature for each wash, and then resuspended in 20 µL water. Bound RNA was released by heating at 75 °C for 5 min and immediately separating beads on a magnetic stand.

### Amplicon-specific library preparation using mutational profiling (MaP)

For reverse transcription, 7 µL of biotin-enriched RNA or 1 µg of total RNA (bulk samples) was combined with 2 µL of 10 mM dNTPs and 1 µL of 2 µM gene-specific primers (10 µL total) [Table S1]. Reverse transcription using SuperScript II under MaP conditions was performed as described previously^54^.

Amplicon libraries were generated using a two-step PCR strategy^62^ (Table S1). For biotin-enriched samples, the entire cDNA product was used as input for the first PCR, whereas 5/12 of cDNA product was used for bulk samples. PCR was performed using Q5 Hot Start High-Fidelity DNA Polymerase (NEB) using the following programs: for AdoCbl, 98°C for 30 s; 20 cycles of 98°C for 10 s, 64°C for 20 s, and 72°C for 20 s; followed by 72°C for 2 min. For 7SK, 98°C for 30 s; 20 cycles of 98°C for 10 s, 66°C for 20 s, and 72°C for 20 s; followed by 72°C for 2 min.

PCR1 products were purified using Mag-Bind TotalPure NGS beads (Omega Bio-tek; 0.8× ratio) and eluted in 12 µL of nuclease-free water. For PCR2, 10–12 µL of PCR1 product was used for biotin-enriched samples, and 2 ng of PCR1 product was used for bulk samples. Reactions were amplified using the program: 98°C for 30 s; 15 cycles of 98°C for 10 s, 65°C for 30 s, and 72°C for 20 s; followed by 72°C for 2 min. PCR2 products were purified with Mag-Bind TotalPure NGS beads (0.7× ratio), assessed for size and quality by TapeStation analysis (Agilent), and sequenced on an Illumina MiSeq instrument using v2 (2 × 150 bp or 2 × 250 bp) or v3 (2 × 300 bp) chemistry.

### Cell-free E. coli rRNA library preparation

Cell-free rRNA libraries were prepared from randomly primed total RNA using nonamers (Sigma) following the Nextera strategy^62^. For reverse transcription, 7 µL of biotin-enriched RNA or 1 µg of total RNA (bulk samples) was combined with 2 µL of 10 mM dNTPs and 1 µL of 200 ng/uL nonamers (10 µL total). Primers were annealed by heating at 65°C for 10 min followed by cooling at 4°C for 2 min. Reactions were then supplemented with MaP buffer containing a custom 10× NTP mix [0.5 M Tris-HCl (pH 8.0), 0.75 M KCl, 0.1 M DTT in 50 µL] and adjusted to 1× [2.2× NTP mix, 2.2 M betaine, 13.3 mM MgCl_2_]. Each reaction received 9 µL of 5× MaP buffer, incubated briefly at room temperature, and initiated with 1 µL of Superscript II reverse transcriptase. Reverse transcription proceeded according to the program: 25°C for 10 min, 42°C for 90 min, 10× (50°C for 2 min and 42°C for 2 min), and 72°C for 10 min. cDNA was purified using SPRI beads (Omega Bio-tek; 1.8× bead ratio). cDNA was converted into double-stranded DNA (dsDNA) using the NEBNext second-strand synthesis module (NEB) using a two hour incubation at 16 °C. dsDNA was purified by SPRI beads (Mag-Bind TotalPure NGS beads; 0.65x ratio). dsDNA was then tagmented with Nextera XT (Illumina) following the manufacturer’s protocol and purified by SPRI beads (Mag-Bind TotalPure NGS beads; 0.56x ratio). Libraries were sequenced on an Illumina MiSeq using 2 x 300 paired-end sequencing.

### In-cell RAID-MaP protocol

RAID-MaP experiments were performed in HEK293T cells expressing NIK3x-GFP-APEX2^1^, LMNA-APEX2^1^, or SRSF1-APEX2^59^. The NIK3x-APEX2 and LMNA-APEX2 constructs were stably expressed under constitutive promoters and localize APEX2 to nucleolar and nuclear lamina environments, respectively. SRSF1-APEX2 cells were a generous gift from B. Blencowe (University of Toronto) and express the APEX2 fusion from a doxycycline-inducible promoter to allow controlled induction prior to labeling. All cell lines were maintained in DMEM (Gibco) with 10% FBS (Gibco), 100 U/mL Pen/Strep (Gibco), 2 mM sodium pyruvate (Gibco), and MEM non-essential amino acids (Gibco) at 37°C and 5% CO2. Cells were regularly tested for and confirmed to be free of mycoplasma.

Cells were plated 24 h prior to probing on 1% fibronectin coated, 10 cm plates to achieve ∼90– 95% confluency at the time of labeling. For SRSF1-APEX2 cells, APEX expression was induced by addition of 2 µg/mL of doxycycline 24 h prior to probing. 30 min prior to probing, cell medium was exchanged with new medium containing 500 µM biotin-phenol. Immediately prior to probing, cells were removed to 23°C for 3 min. 23°C was then used as the labeling temperature to limit RNA diffusion, consistent with established APEX protocol^1^. DMS labeling was initiated by replacing medium with 3 mL of pre-mixed DMS probing medium. Probing medium was prepared immediately prior to DMS reactions by combining 2.34 mL of fresh medium, 200 mM of bicine buffer (pH 8.0), and 60 µL of 100% DMS and mixing vigorously to ensure homogeneity. Cells were incubated at 23 °C for 2 min with occasional gentle agitation. APEX labeling was then initiated by adding 36 µL of 98 mM H_2_O_2_ (1 mM final conc.), gently agitating the cells, and incubating DMS-H_2_O_2_ treated cells for an additional 1 min, resulting in a total DMS treatment time of 3 min and H_2_O_2_-treatment time of 1 min. Reactions were quenched by aspirating the medium followed by adding 2 mL of 10% β-mercaptoethanol BME in PBS. APEX-only controls were processed in parallel as previously described^1,63^, with quenching performed using 10% BME in PBS. Unlabeled controls were processed identically except for omission of hydrogen peroxide.

Cells were immediately scraped, collected by centrifugation at 4,000 rpm for 5 min at 4°C, and resuspended in lysis buffer (Buffer RLT Plus, RNeasy Plus Mini Kit, Qiagen). Lysates were either processed immediately or stored at 4°C for several hours. Lysates were passed through genomic DNA removal columns (RNeasy Plus Mini Kit, Qiagen) and RNA purified according to the manufacturer’s protocol, substituting Buffer RW1 with Buffer RWT. RNA concentration was determined by NanoDrop, and integrity was assessed using RNA TapeStation (Agilent). Only samples with RIN scores >8.5 were used for downstream analyses. For all experiments, data was collected in triplicates for every sample condition.

### Biotin enrichment of in-cell probed RNA

RNA from in-cell probing experiments was biotin-enriched as described^63^ using Pierce streptavidin magnetic beads (ThermoFisher). For DMS+APEX treated RNA, ∼45 µg of RNA was used per enrichment, and ∼30 µg of RNA per enrichment for APEX-only treated samples. 10 μl beads were used per 25 µg of RNA. The beads were washed 3 times in B&W buffer (5 mM Tris-HCl pH 7.5, 0.5 mM EDTA, 1 M NaCl, 0.1% TWEEN 20 (Sigma Aldrich)), followed by 2 washes in Solution A (0.1 M NaOH and 0.05 M NaCl), and 1 wash in Solution B (0.1 M NaCl). The beads were then suspended in 150 μL 0.1 M NaCl and incubated with ∼125 μL RNA diluted in water on a rotator for 2 h at 4°C. The beads were then placed on a magnet and the supernatant discarded. Beads were washed 3 times in B&W buffer and resuspended in 54 μL water.

RNA was released by treating beads with 2 mg/mL Proteinase K (Ambion) in proteinase digestion buffer (1x: 330 μL 10X PBS (pH 7.4), 330 μL 6.6% N-Lauryl sarcosine sodium solution (Sigma Aldrich), 66 μL 165 mM EDTA, 16.5 μL 330 mM DTT, 3 μL Ribolock RNase inhibitor, 357.5 H_2_O). The beads were then incubated at 42°C for 1 h, followed by 55°C for 1 h on a shaker. The RNA was then purified using RNA clean and concentrator columns (Zymo). Enriched RNA was either eluted into 5 μL H_2_O for RNA-seq/rRNA library preparation or up to 14 μL for samples used in qPCR and amplicon library preparation.

### Dot blot analysis

Dot blots were performed on biotinylated RNA as described previously^63^. Briefly, 1-2 μg of purified RNA was blotted on an Amersham Protran 0.45 nitrocellulose membrane, and the membrane was incubated at 37°C for 15 min to allow liquid to dry. The RNA was crosslinked to the membrane using 2500 μJ energy (254 nm wavelength, Stratalinker 2400). The membrane was then incubated with 15 mL PBST containing 1 μL LI-COR Streptavidin IRDye 800CW, washed three times with PBS, and imaged on an Amersham Typhoon 5 (Cytiva). After imaging for biotin, the membrane was incubated with methylene blue solution (0.02% methylene blue with 0.3M sodium acetate) for ∼10 minutes before being washed with PBS and imaged again. Only samples showing strong blot intensity were selected for further experiments.

### Sequencing library preparation for nucleolar RAID-MaP experiments

Total RNA libraries were prepared from biotin-enriched RNA after RAID-MaP in NIK-APEX2 cells (corresponding to ∼30–45 µg of pre-enriched RNA). For biotin-enriched samples, 5 µL (entire yield after biotin enrichment) was used for library preparation, while 1 µg of unenriched/bulk RNA was used in parallel. Input RNA was fragmented by incubation with 13 µL of Fragment, Prime, Finish (FPF) mix from TruSeq Stranded mRNA Library Preparation Kit (Illumina) with the listed modifications at 94°C for 6 min, followed by rapid cooling to 4°C to yield fragments of ∼210 nt.

Reverse transcription was carried out in SSII-MaP conditions^62^, containing 50 mM Tris-HCl (pH 8.0), 75 mM KCl, 10 mM DTT, 6 mM MnCl_2_, 1 M betaine, 0.3 mM dNTPs, 5 ng/µL random nonamers, and 1.4 U Superscript II (Invitrogen). Reactions were incubated at 25°C for 10 min, 42°C for 90 min, 10× (50°C for 2 min, 42°C for 2 min), and finally 72°C for 10 min. cDNA was purified using SPRI beads (Mag-Bind TotalPure NGS, Omega Bio-tek; 1.8× ratio) and eluted in 32 µL nuclease-free water. Second-strand synthesis was performed by incubating the purified cDNA with 20 µL of Second Strand Marking Master Mix (Illumina) for 1 h at 16°C. DNA was again purified with SPRI beads (1.8× ratio) and used as input for sequencing library preparation following the manufacturer’s protocol.

Final libraries were assessed for size distribution and quality using a TapeStation system (Agilent) with High Sensitivity ScreenTape. Libraries were sequenced using paired-end 2×75 bp reads on either a NovaSeq or MiSeq platform (Illumina).

### Actinomycin D timecourse experiments

HEK293T-NIK3x-GFP-APEX2 cells were grown in DMEM on 1% fibronectin coated plates for 24 h till 90-95% confluency before experiment. The media was then exchanged DMEM supplemented with 10 ng/mL actinomycin D (Sigma) at 37°C for 1, 2, or 3 h. During the final 30 min of incubation, medium was replaced with fresh medium containing 10 ng/mL actinomycin D and 500 µM biotin-phenol. Subsequent steps were performed according to the general RAID-MaP protocol (see above). Biological replicates (n = 3) were collected on independent days. For the 3 h timepoint, one of the 3 h replicates failed post-enrichment QC and was thus discarded.

### RT–qPCR validation of nuclear RNA enrichment

Enrichment of nuclear-specific RNA candidates in HEK cells expressing LMNA-APEX2 and SRSF1-APEX2 cells was evaluated by RT–qPCR against *MALAT1* (nuclear), and negative controls were *GAPDH* and *MTCO2* (Table S1). Enriched RNA was reverse transcribed using the MaP protocol (described above) with 1 µL random hexamer primers (ThermoFisher). qPCR was performed with the indicated primer sets using 2× SYBR Green PCR Master Mix (ThermoFisher Scientific) on a LightCycler 480 system (Roche). Relative enrichment was calculated as the ratio of transcript abundance in labeled samples relative to bulk controls, normalized to *GAPDH* using the ΔΔCt method^64^. Samples showing no relative enrichment were not used for downstream library preparation.

### rRNA read-depth analysis

Sequencing data from nucleolar in-cell samples were processed using ShapeMapper v2.2 with the –dms flag and aligned to RN45S5 (NR_046235). Processed ShapeMapper reactivity.txt files were then analyzed using custom Python scripts. To account for differences in sequencing depth between libraries, read depths were normalized by dividing the read depth at each nucleotide by the average read depth across the entire sample. Normalized read depths corresponding to the 12,993 nucleotides of the human pre-rRNA transcript were extracted, encompassing the 5′ external transcribed spacer (5′ETS), 18S rRNA, internal transcribed spacer 1 (ITS1), 5.8S rRNA, internal transcribed spacer 2 (ITS2), and 28S rRNA regions. Normalized read depths were summed across defined rRNA processing segments and divided by the total read depth to calculate the percentage of signal associated with each region for each sample.

### rRNA cleavage junction analysis

Alignment data were processed from SAM files to assess read coverage across known rRNA cleavage junctions. SAM files were parsed using a custom Python script to count reads spanning approximate known rRNA cleavage junctions (positions 422, 1643, 3655, 5551, 6157, 6476, 6947, 7926), including a ±5 nucleotide window around each site to ensure the site was included in the read. The number of reads spanning each junction was then normalized by dividing by the total number of reads in the sample. Enrichment was then obtained by dividing the normalized count from nucleolar samples by the normalized count from bulk samples.

### RAID-MaP data processing and normalization

Sequencing data from all RAID-MaP samples were processed using ShapeMapper v2.2 with the --dms flag^29^. The 18S rRNA gene sequences were aligned Rfam RF01960, 28S rRNA sequences were aligned to Rfam RF02543, and 7SK were aligned to RF00100. 7SK RAID-MaP libraries were prepared using unique molecular barcodes (UMIs), and fastq files were pre-processed to remove duplicate reads using UMICollapse^65^ via the automated pipeline https://github.com/MustoeLab/umi_dedup_pipeline^60^.

Per-nucleotide mutation rates were extracted from ShapeMapper2 output profiles, requiring a minimum read depth of 500 at each position (depthcut = 500). Background subtraction was performed by computing the mean mutation rate across background replicates (either untreated for bulk samples, or APEX only-enriched for RAID-MaP samples) and subtracting this mean from the mutation rate measured in each DMS treated sample. Each pair of unenriched and enriched samples was normalized together by aggregating the background subtracted A and C reactivities from both samples and then dividing by the average reactivity of the 90th-99th percentile most highly reactive nucleotides^66^. For rRNA datasets, 18S and 28S rRNAs were also aggregated together during the normalization process. Normalization was performed independently for replicate. G and U positions were excluded from quantitative comparisons due to low modification rates under our DMS probing conditions.

Differential reactivity, ΔR, between enriched and unenriched conditions was computed at each A or C nucleotide as ΔR = R(enriched) − R(unenriched), where reactivity values below zero were rounded to zero prior to subtraction.

For sliding-window analysis (window of 3 nucleotides, centered on each position), average nucleotide reactivity was calculated using a sliding window spanning positions n–1, n, and n+1. Normalized reactivity were averaged across the three positions and four biological replicates (12 measurements total) to obtain a regional mean, and assessed for statistical significance using a Wilcoxon signed-rank test. Differences with |ΔR| < 0.01 were assigned p = 1 regardless of the statistical test result. Positions reaching p < 0.05 were counted as significantly differentially reactive. All replicates verified to be enriched for the compartment of interest were used in the analysis [n=4 for NIK3x-APEX2, n=3 for SRSF1-APEX2, n=2 for LMNA-APEX2].

### Proximal protein and nucleotide analysis

Spatial proximity analysis between rRNA and proteins or other rRNA nucleotides was performed using PyMOL (v3.0)^67^. The mature rRNA cryo-EM structure (PDB: 6QZP) was used for analysis. All resolved nucleotide positions in the both rRNA subunits were enumerated. For each nucleotide, protein or nucleotide residues within a 5 Å cutoff were identified using a distance-based selection criterion in PyMOL.

### 7SK FISH analysis

Approximately 10,000 cells per well were plated in 1% fibronectin-coated 8-well chamber slides and grown for 48 hours to reach ∼80–90% confluency at the time of experiment. For SRSF1-APEX2 cells, APEX2 expression was induced by addition of 2 µg/mL doxycycline in culture medium 24 hours after plating, at approximately 40% confluency.

For immunofluorescence, cells were washed once with ice-cold PBS supplemented with Ca^2+^ and Mg^2+^, then fixed with 4% paraformaldehyde (PFA) in RNase-free PBS on ice for 30 minutes. Fixative was removed and cells were washed once with PBS at room temperature. Cells were permeabilized with 0.5% Triton X-100 in PBS for 30 minutes at room temperature, followed by two washes, 3 minutes each with PBS. Primary antibody (anti-FLAG rat antibody, 1:500 in 2× SSC buffer) for SRSF1 was applied and incubated at 37°C on a rotator for 1 hour. Cells were washed three times with 2× SSC buffer, then incubated with secondary antibody (anti-rat Alexa Fluor 488, 1:500 in 2× SSC buffer) for 30 minutes at room temperature. Cells were washed four times with 2× SSC buffer, then post-fixed with 4% PFA in RNase-free PBS for 15 minutes at room temperature. Fixative was removed and cells were washed once with PBS (∼5 minutes).

For FISH, cells were briefly pre-washed once for 2 minutes in wash buffer A (2× SSC, 10% formamide in nuclease-free water). Hybridization was performed in buffer containing 2× SSC, 10% formamide, and 10% dextran sulfate in nuclease-free water, supplemented with 7SK FISH probes(IDT) at 1:500 dilution. Samples were incubated at 37°C overnight in a humidity chamber. Following hybridization, probe solution was removed and cells were washed once in wash buffer A for 30 minutes at 37°C, followed by a second wash in wash buffer A containing DAPI (1 µg/mL) for 30 minutes at 37°C. Cells were then washed once in wash buffer B (2× SSC in nuclease-free water) for 5 minutes at room temperature. Samples were imaged in 2× SSC buffer. Samples were imaged on the iSIM as listed above.

### 7SK structural population deconvolution

Bulk 7SK samples were deconvolved using the using DanceMapper pipeline^54^. Data from the three SRSF1-APEX replicates were merged together and then deconvolved using DanceMapper using -maskU and -maskG flags. Enriched (compartment-specific) samples had insufficient numbers of unique reads after deduplication to be able to deconvolve. Thus, we estimated (A, B, H) state populations by decomposing the measured mean reactivity into a linear combination of the three state reactivity profiles. Specifically, we solved the system **r** = **Sp** using the Moore-Penrose pseudoinverse (numpy.linalg.lstsq, rcond=None), where **r** is the per-nucleotide reactivity vector for a given enriched sample, **S** is the matrix of state reactivities determined from bulk DANCE analysis (nucleotides × 3 states), and **p** is the vector of state population weights. Only nucleotides with non-missing values across all three states and the target compartment were included. The resulting population weights were normalized to sum to one to yield fractional state populations.

### Immunofluorescence staining and microscopy

Cells were seeded on 10 cm dishes containing coverslips coated with 1% fibronectin and cultured until ∼90% confluence. For DMS-treated cells, medium was exchanged with DMS-containing medium and incubated for 3 min at 23°C. Coverslips from either untreated or DMS treated cells were then retrieved, excess medium removed, and cells fixed in 4% paraformaldehyde in PBS for 15 min at 23°C. Following fixation, cells were washed three times with PBS and permeabilized in 0.05% Triton X-100 in PBS for 15 min. Cells were washed again three times with PBS and blocked for 1 h at room temperature in antibody diluent (ThermoFisher). Cells were incubated overnight at 4 °C with primary antibodies diluted in blocking buffer with their respective antibodies. To visualize APEX fusions, LMNA-APEX2 and SRSF1-APEX2 were stained with rabbit anti-V5 (Cell Signaling Technology, cat. 13202T, 1:500). SRSF1-APEX2 cells were also co-stained with mouse anti-SC35 as nuclear speckle marker (Abcam, cat. ab11826, 1:500). NIK3x-GFP-APEX2 cells were stained with nucleolar DFC marker anti-Fibrillarin (Abcam, cat. ab5821). After three washes in PBST (PBS with 0.05% Tween-20), coverslips were incubated with secondary antibodies for 1 h at room temperature: Alexa Fluor 488 (ThermoFisher, cat. A-11008, 1:2000) for APEX, Alexa Fluor 568 (ThermoFisher, cat. A-11004, 1:2000) for compartment markers. After three washes in PBST, cells were mounted on coverslips using DAPI mounting media (VWR).

Confocal imaging was performed using a Zeiss LSM 900 microscope (Axio Observer.Z1/7) equipped with an Airyscan 2 detector and a Plan-Apochromat 63×/1.40 NA oil immersion objective. Images were acquired with 2.0× digital zoom at a resolution of 1413 × 1413 pixels, corresponding to a pixel size of 0.035 µm. Imaging was performed in frame scan mode with bidirectional scanning, a scan speed of 6, and a pixel dwell time of 1.47 µs. Line averaging of 16 was applied. The pinhole was set to 5.0 Airy Units (248 µm). Excitation was performed using appropriate laser lines (405, 488, 561, or 640 nm) depending on the fluorophore used. Emission was collected over a detection range of 450–700 nm using a GaAsP-PMT detector. For each channel, laser power and detector gain were optimized to avoid saturation and were kept constant across all conditions within each experiment. Laser blanking was enabled. Airyscan super-resolution imaging was performed in 2D SuperResolution mode (automatic processing strength: 5.8). Raw images were processed using Zeiss ZEN software with identical Airyscan processing parameters applied across all samples. All images were acquired as single optical sections under identical acquisition settings within each experiment. Comparisons of fluorescence intensity were performed only between images acquired under identical settings.

### iSIM microscopy for Actinomycin D time course

HEK293T-NIK3x-GFP-APEX2 cells were plated onto 1% fibronectin–coated 8-well chamber slides (Ibidi) at ∼50% confluency. Prior to imaging, cells were treated with culture medium containing actinomycin D at final concentration of 10 ng/mL, or with vehicle control (DMSO) or left untreated. Imaging was performed using a microscope system consisting of a Nikon Ti2-E inverted microscope equipped with a VisiTech instant SIM (VT-iSIM), a Hamamatsu ORCA-Quest qCMOS camera, and a CFI60 Plan Apochromat Lambda D 100× oil immersion objective. Data was collected at 20-min intervals for 3.5 h. Cells were maintained at 37°C with 5% CO_2_ during imaging.

### ASO treatment and imaging in U2OS cells

U2OS cells expressing endogenously tagged NPM1-mCherry and Pol1-muGFP^51^ were cultured in sterile-filtered McCoy’s 5A Modified Medium (1×) with L-glutamine (Gibco) supplemented with 10% fetal bovine serum (FBS) and 1% penicillin–streptomycin (Gibco). Cells were plated in 8-well imaging chambers (Ibidi) and grown until ∼80% confluency. All cultures were tested for mycoplasma contamination every two weeks using a Mycoplasma Test Kit (Applied Biological Materials) following the manufacturer’s protocol.

Cells were transfected with antisense oligonucleotides (ASOs; Table S1) using Lipofectamine RNAiMAX (ThermoFisher). For each well, transfection complexes were prepared in 25 µL Opti-MEM containing 0.75 µL RNAiMAX and 36 pmol ASO. Prior to transfection, the culture medium was replaced with Opti-MEM. Cells were incubated with the transfection mixture for 6 hours, after which the medium was replaced with standard U2OS growth medium (McCoy’s 5A supplemented with 10% FBS and 1% penicillin–streptomycin). In addition to experimental wells, two control conditions were included: a 'Control' condition receiving neither ASO nor transfection reagent, and a 'Mock' condition receiving transfection reagent alone.

Twenty-four hours after transfection, cells were imaged for NPM1-mCherry and Pol1-muGFP. Cells were then washed three times with warm PBS and fixed with 4% paraformaldehyde (PFA) for 10 minutes at room temperature. Following fixation, cells were washed three times with PBS. ASOs were labeled using the Click-iT RNA Imaging Kit protocol (ThermoFisher). Cells were permeabilized with 0.5% Triton X-100 in PBS (300 µL per well) for 15 minutes at room temperature with gentle shaking. Cells were then washed with PBS and incubated with 125 µL of the Click-iT reaction cocktail for 30 minutes at room temperature with gentle shaking, protected from light. The cells were then washed once with Click-iT reaction rinse buffer followed by additional PBS rinse.

Imaging was performed using a microscope system consisting of a Nikon Ti2-E inverted microscope equipped with a VisiTech instant SIM (VT-iSIM), a Hamamatsu ORCA-Quest qCMOS camera, and a CFI60 Plan Apochromat Lambda D 100× oil immersion objective. Sequential excitation was used for each fluorophore: Alexa Fluor 647 (640 nm), NPM1-mCherry (561 nm), and Pol1-muGFP (488 nm). No detectable spectral bleed-through was observed between channels.

Approximately every other day, power meter measurements were acquired at the focal plane to verify microscope performance and enable day-to-day comparison of measurements. Fluorescent dye standards were used to calibrate the digital level (DL) offset and DL-to-photon conversion. Calibration was performed by fitting the histogram of photon counts at low illumination levels to determine the DL offset and conversion factor. These measurements were used to generate a conversion table referenced to acquisition settings, as well as background and flat-field correction functions (implemented as functions because they depended on laser intensity). Camera linearity was verified manually as needed.

### Quantification of NPM1 concentration

Imaging data were analyzed in Micro-Manager using a custom-built imaging plugin adapted from AcqEngJ and NDViewer. For all images, background subtraction, flat-field correction, and DL-to-photon conversion were applied prior to quantification. During intensity quantification within regions of interest (ROIs), the average intensity within each ROI was calculated on the background-subtracted image and the flat-field correction was applied separately; corrected intensity values were then obtained by dividing the background-subtracted signal by the corresponding flat-field values.

ROIs corresponding to nuclei were identified based on endogenous NPM1–mCherry fluorescence (561 nm channel) using a combination of thresholding and Gaussian blur. Segmented nuclei were manually inspected and corrected when necessary. In cases where automated threshold-based segmentation was insufficient, nuclei were manually segmented using circular ROIs. Representative regions within the nucleoplasm were manually selected on a per-cell basis. Representative regions of nucleoli were identified by determining the local intensity maximum within nuclear ROIs using a sliding box of 6 × 6 pixels. Analysis of relative NPM1 concentrations was done in images of live cells.

### Kinetic modeling of rRNA structural changes after ActD treatment

Time-resolved changes in ΔRwere modeled using a boundary-constrained exponential function to estimate domain-level folding kinetics. For each nucleotide, ΔReactivity measurements from all biological replicates across four timepoints (0, 1, 2, and 3 h following ActD treatment) were fit to the model:

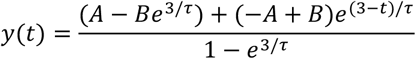

Where *t* is time, *A* and *B* represent the fitted ΔReactivity values at the initial and final boundaries per nucleotide, and *τ* represents the characteristic time constant describing the structural change. Model fitting was performed using non-linear least squares optimization implemented in SciPy (curve_fit). All replicate measurements were included in the fit, excepting nucleotides that lacked measurements at either the 0 h or 3 h endpoints and therefore have undetermined boundaries. Nucleotides were first fit individually using the median ΔReactivity values observed at the 0 h and 3 h timepoints as the initial parameter estimates for A and B, respectively, the constraint 0 < *τ* < 30 h. Nucleotide-level fits were subsequently filtered to retain only well-behaved trajectories (0.1 < *τ* < 10 h). Domain-level kinetics were then estimated by fitting all nucleotides within each rRNA domain simultaneously using a shared *τ* parameter while allowing nucleotide-specific *A* and *B* fitted values, using A and B parameters from nucleotide-level fits as initial parameter guesses. This global fitting approach captures a common kinetic timescale for structural transitions within each domain that is more robust to noise present in nucleotide-level fits. Uncertainty in the fitted time constant (*τ*) was estimated from the covariance matrix returned by the non-linear least squares optimization.

Domain-level kinetic trajectories were visualized by evaluating the fitted exponential model using the domain-shared *τ* and the median of nucleotide-specific A and B parameters. To enable cross-domain comparison, trajectories were normalized by shifting to zero at t = 0 and scaling by the factor (1 − e^(−3/*τ*)^) / (B − A), then inverted so that decreasing ΔR appears as positive folding progression.

