## Supplementary Figures and Tables for "Mapping RNA structure assembly and remodeling in biomolecular condensates"

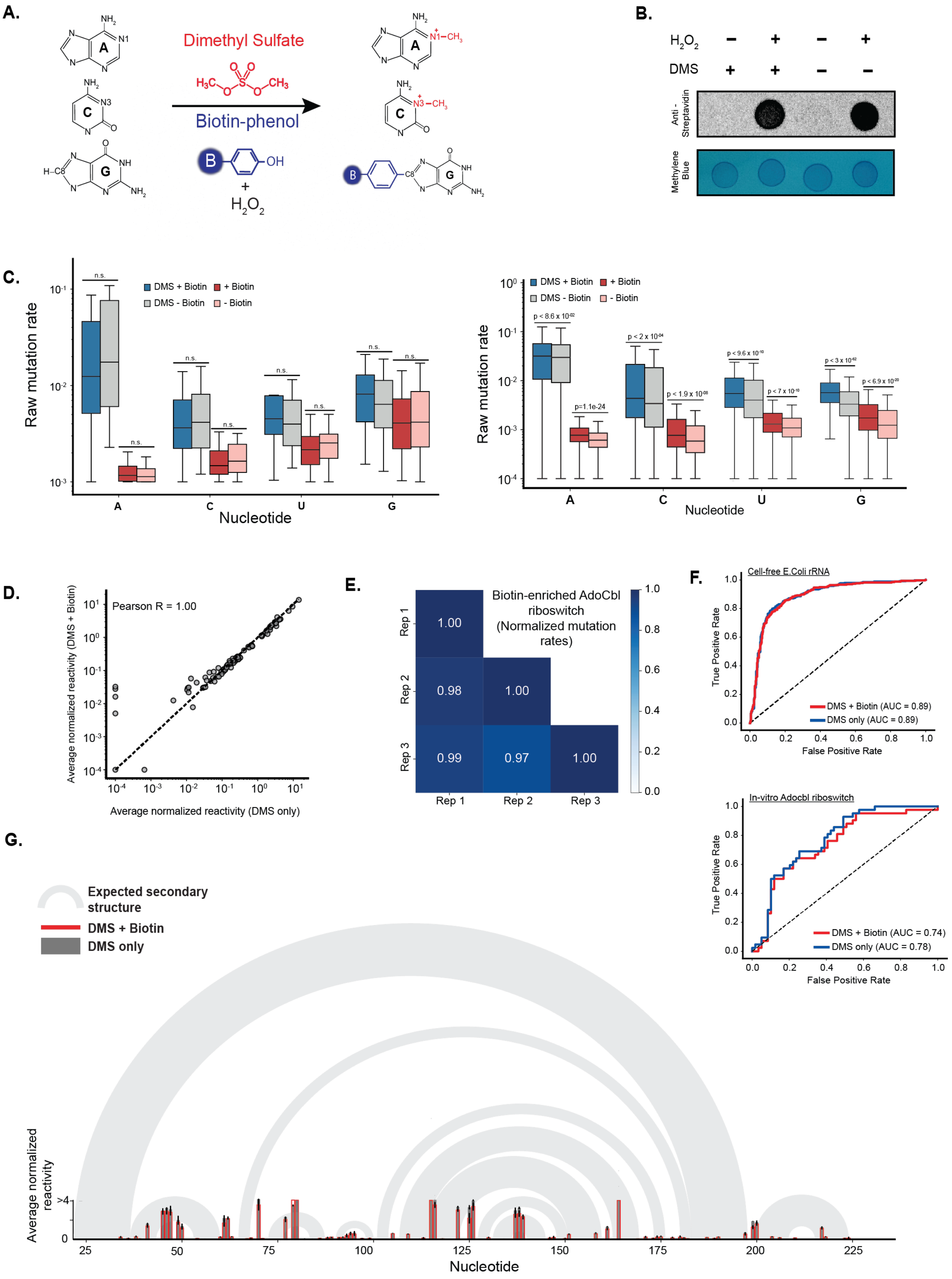
A.
Dimethyl Sulfate
Biotin-phenol
+
H2O2
B.
H2O2
DMS
Anti - Streptavidin
Methylene Blue
C.
Raw mutation rate
DMS + Biotin
+ Biotin
DMS - Biotin
- Biotin
n.s.
Nucleotide
p=1.1e-24
D.
Pearson R = 1.00
Average normalized reactivity (DMS + Biotin)
Average normalized reactivity (DMS only)
E.
Biotin-enriched AdoCbl riboswitch (Normalized mutation rates)
Rep 1
Rep 2
Rep 3
1.00
0.98
1.00
0.99
0.97
1.00
F.
Cell-free E.Coli rRNA
True Positive Rate
False Positive Rate
DMS + Biotin (AUC = 0.89)
DMS only (AUC = 0.89)
In-vitro Adocbl riboswitch
DMS + Biotin (AUC = 0.74)
DMS only (AUC = 0.78)
G.
Expected secondary structure
DMS + Biotin
DMS only
Average normalized reactivity
Nucleotide

**Figure S1. In vitro validation of RAID-MaP on model RNAs.**

- **A.** Chemical adducts generated by dimethyl sulfate (DMS) and radical-based biotinylation.
- **B.** Dot blot confirming RNA biotinylation. E. coli rRNA was treated with DMS (6 min, 37°C) followed by a 1-min APEX-mediated biotinylation reaction (HRP, $H_2O_2$).
- **C.** Box plots showing measured mutation rates at all four nucleotide positions in untreated, DMS-only, and DMS+APEX samples. Mann–Whitney U test for AdoCbl riboswitch (left) and Cell-free E.Coli rRNA (right).
- **D.** Correlation of adenosine and cytosine reactivities between DMS-only and DMS+APEX AdoCbl RNA samples (R > 0.95).
- **E.** Pearson R between normalized DMS reactivities measured from biotin-enriched RNA from three indepedent replicate experiments performed on the AdoCbl riboswitch.
- **F.** Receiver operating characteristic (ROC) analysis comparing area under the curve (AUC) values for DMS-only and DMS+APEX samples against the expected reference secondary structure for cell-free probed *E. coli* rRNA and AdoCbl riboswitch.
- **G.** DMS-MaP profiles recapitulate known RNA secondary structure after biotinylation. Arc diagram of AdoCbl RNA showing cryo-EM-derived base pairs (gray arcs) overlaid on RAID-MaP reactivity.

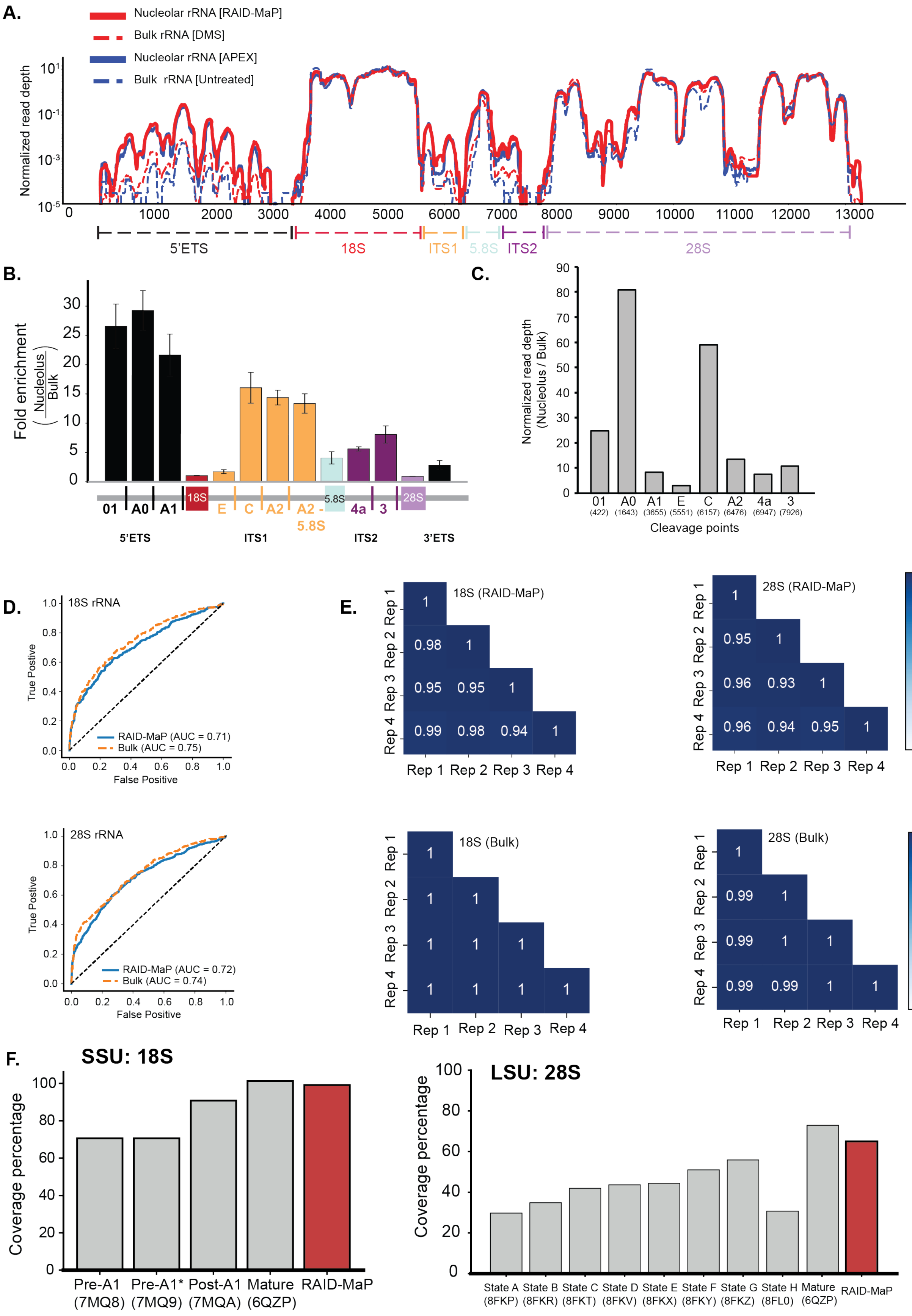
A.
Nucleolar rRNA [RAID-MaP]
Bulk rRNA [DMS]
Nucleolar rRNA [APEX]
Bulk rRNA [Untreated]
Normalized read depth
5'ETS
18S
ITS1
5.8S
ITS2
28S
B.
Fold enrichment (Nucleolus / Bulk)
01
A0
A1
E
C
A2
A2 - 5.8S
4a
3
5'ETS
ITS1
ITS2
3'ETS
C.
Normalized read depth (Nucleolus / Bulk)
Cleavage points
D.
18S rRNA
True Postive
False Positive
RAID-MaP (AUC = 0.71)
Bulk (AUC = 0.75)
28S rRNA
RAID-MaP (AUC = 0.72)
Bulk (AUC = 0.74)
E.
18S (RAID-MaP)
28S (RAID-MaP)
18S (Bulk)
28S (Bulk)
Rep 1
Rep 2
Rep 3
Rep 4
F.
SSU: 18S
Coverage percentage
Pre-A1 (7MQ8)
Pre-A1* (7MQ9)
Post-A1 (7MQA)
Mature (6QZP)
RAID-MaP
LSU: 28S
State A (8FKP)
State B (8FKR)
State C (8FKT)
State D (8FKV)
State E (8FKX)
State F (8FKY)
State G (8FKZ)
State H (8FL0)
Mature (6QZP)
RAID-MaP

**Figure S2. RAID-MaP produces structural maps specific to GC.**

- **A.** Read depth across the 47S precursor in bulk and RAID-MaP samples. Per-nucleotide read depth normalized to total read depth is shown for bulk (DMS-modified, pre-enrichment) and GC-enriched (biotin-enriched) samples across each processing segment.
- **B.** Mean enrichment of 47S precusor segments. Values were averaged across nucleotides within each segment and across replicates. Error bars denote standard errors.
- **C.** Fold enrichment of cleavage junctions in GC compared to bulk measurements in the 47S rRNA.
- **D.** ROC analysis comparing AUC values for DMS-only and DMS+APEX samples against reference secondary structures.
- **E.** Heatmaps of pairwise Pearson correlations across four replicates for 18S and 28S rRNA under bulk and enriched conditions.
- **F.** RAID-MaP extends nucleotide coverage relative to available cryo-EM structures. Bar plots comparing the fraction of 18S and 28S rRNA nucleotides with structural data in RAID-MaP datasets versus existing cryo-EM models.

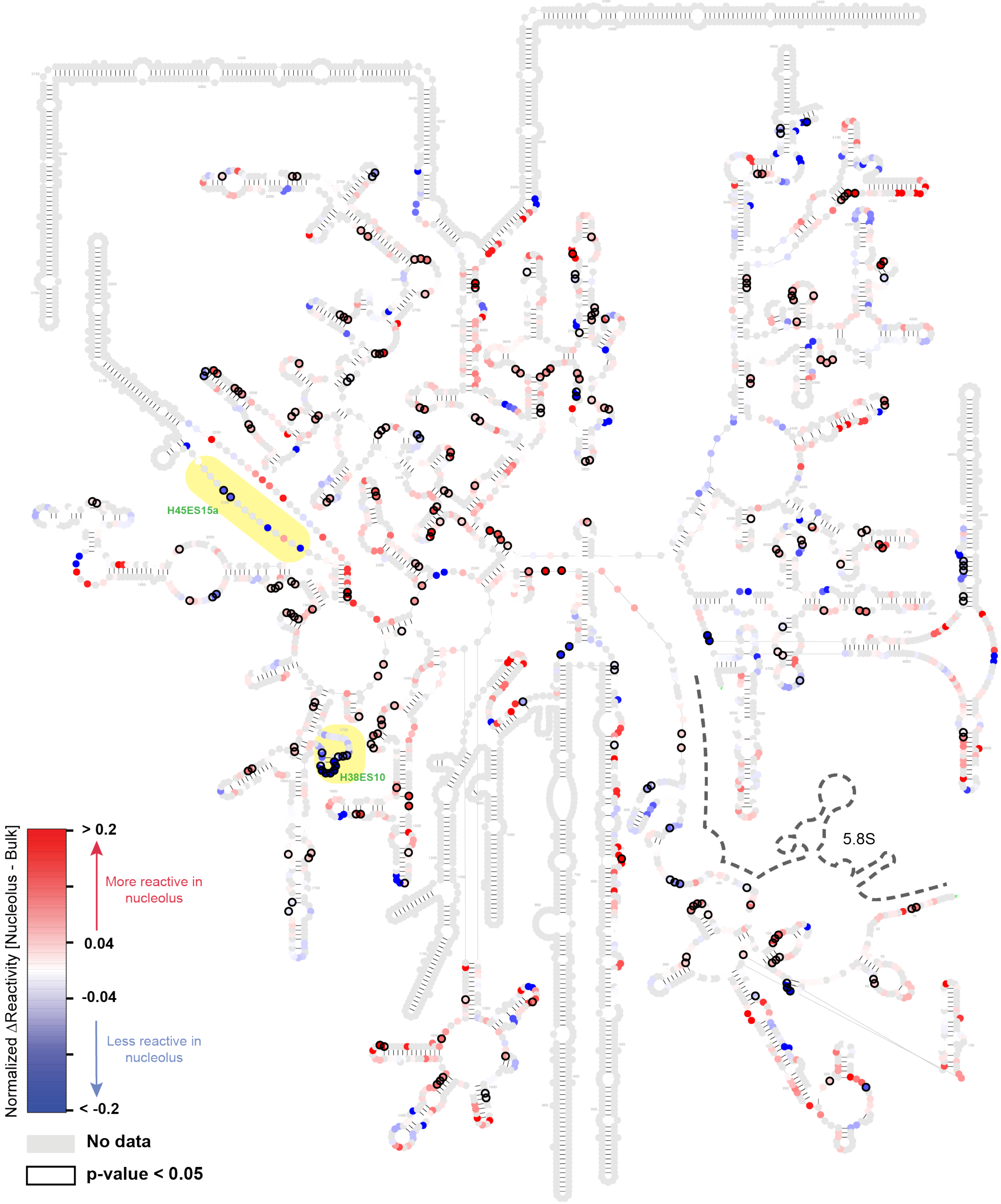
H45ES15a
H38ES10
5.8S
Normalized ΔReactivity [Nucleolus - Bulk]
> 0.2
More reactive in nucleolus
0.04
-0.04
Less reactive in nucleolus
< -0.2
No data
p-value < 0.05

**Figure S3. Per nucleotide structural map of 28S rRNA in the nucleolus.**

Window-averaged ΔR across the 28S rRNA. Regions with increased (red) or decreased (blue) reactivity averaged over a window of 3 nucleotides are plotted on a symlog scale. Black outlined circles denote nucleotides with statistically significant differences ($p < 0.05$, Wilcoxon rank-sum test). Nucleotides involved in 18S–28S subunit interfaces are highlighted in purple, and those participating in long-range tertiary contacts are shown in green.

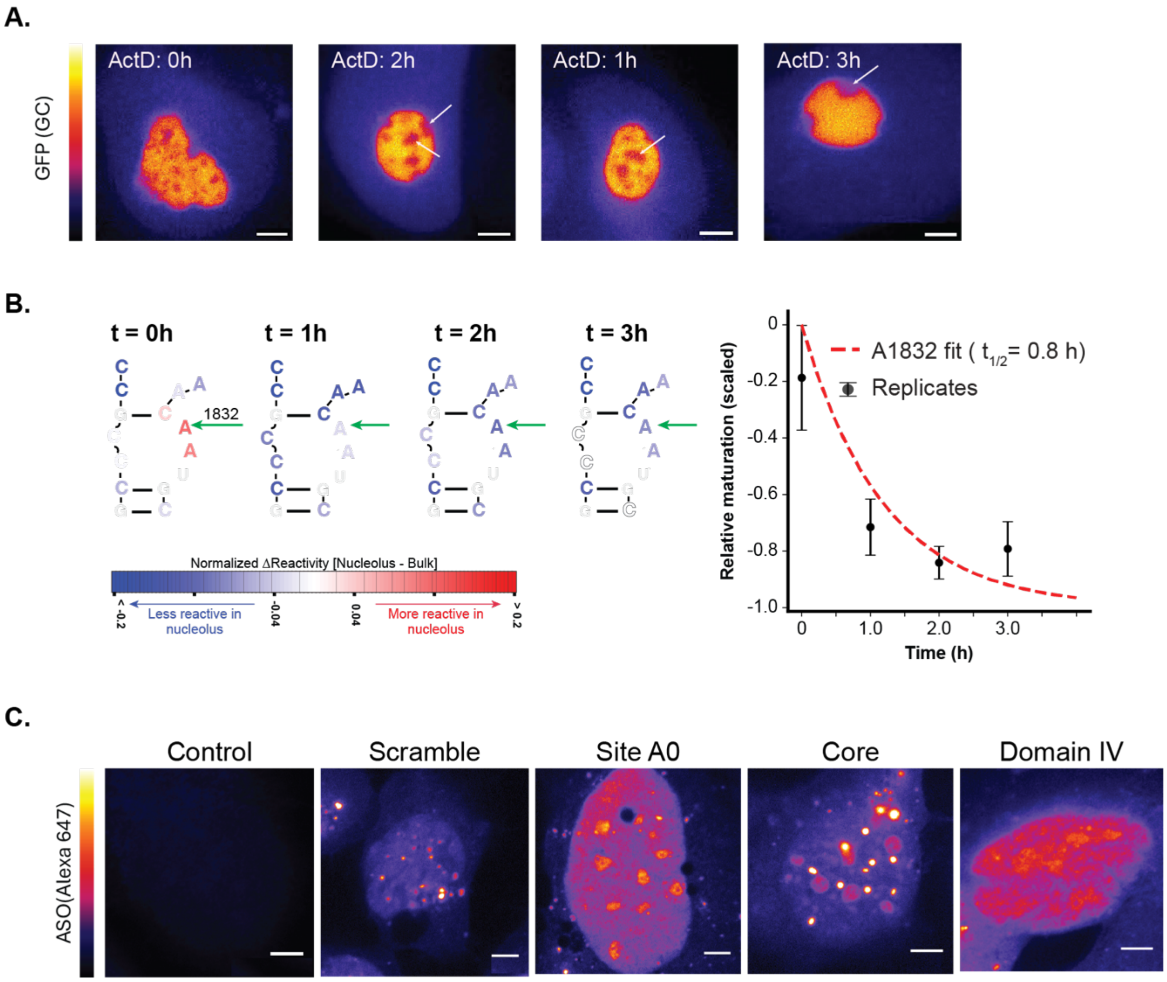
A.
GFP (GC)
ActD: 0h
ActD: 2h
ActD: 1h
ActD: 3h
B.
t = 0h
t = 1h
t = 2h
t = 3h
1832
Normalized ΔReactivity [Nucleolus - Bulk]
< -0.2
-0.04
0.04
> 0.2
Less reactive in nucleolus
More reactive in nucleolus
A1832 fit ( t1/2= 0.8 h)
Replicates
Relative maturation (scaled)
0
-0.2
-0.4
-0.6
-0.8
-1.0
0
1.0
2.0
3.0
Time (h)
C.
ASO(Alexa 647)
Control
Scramble
Site A0
Core
Domain IV

**Figure S4. Kinetic measurements of rRNA structural maturation following transcriptional arrest with actinomycin D.**

- **A.** Confocal images of GC-targeted NIK3x-GFP-APEX expressing cells at 1, 2, and 3 h post-ActD treatment show displacement of FC/DFC components (gaps in GFP signal; white arrows) while the GC remains intact but reduced in size. Scale bar, 2 µm.
- **B.** Per-nucleotide kinetic modeling of structural consolidation. Normalized ΔR at a representative 18S rRNA nucleotide (A1832) across the ActD time course, fit to a boundary-constrained single-exponential model (Methods). Nucleotides are colored by ΔR at each timepoint (red, more reactive in nucleolus; blue, less reactive in nucleolus) on the 18S secondary structure. Green arrows indicate the representative nucleotide A1832 ($t_{1/2}$ = 0.8 h). Points represent means across biological replicates; error bars denote standard error of the mean.
- **C.** Nucleolar localization of fluorescent ASOs. Confocal images of HEK293T cells transfected with fluorescently labeled ASOs (fire LUT, Alexa 647). Scale bar, 4 µm.

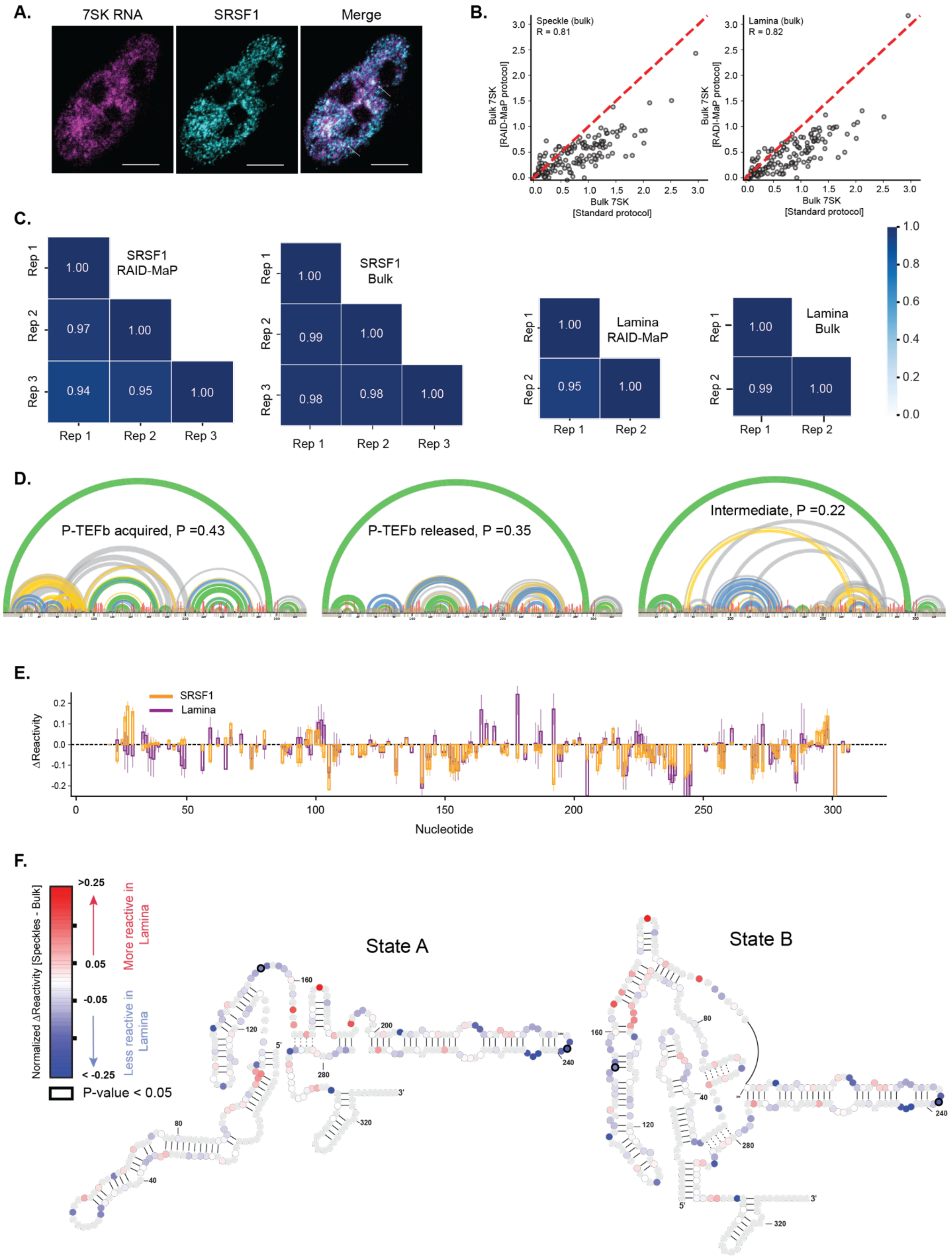
A.
7SK RNA
SRSF1
Merge
B.
Speckle (bulk)
R = 0.81
Bulk 7SK [RAID-MaP protocol]
Bulk 7SK [Standard protocol]
Lamina (bulk)
R = 0.82
Bulk 7SK [RADI-MaP protocol]
Bulk 7SK [Standard protocol]
C.
SRSF1 RAID-MaP
SRSF1 Bulk
Lamina RAID-MaP
Lamina Bulk
Rep 1
Rep 2
Rep 3
D.
P-TEFb acquired, P =0.43
P-TEFb released, P =0.35
Intermediate, P =0.22
E.
SRSF1
Lamina
ΔReactivity
Nucleotide
F.
Normalized ΔReactivity [Speckles - Bulk]
>0.25
0.05
-0.05
< -0.25
More reactive in Lamina
Less reactive in Lamina
P-value < 0.05
State A
State B

**Figure S5. 7SK structural remodeling in nuclear speckles and lamina.**

- **A.** FISH imaging of 7SK RNA (Cy3, magenta) in SRSF1-APEX2 cells co-stained with anti-FLAG antibody to mark nuclear speckles (SRSF1, teal). The merged image shows partial colocalization of 7SK with SRSF1-positive speckles; arrows indicate representative overlap foci. Scale bar, 5 µm.
- **B.** Correlation between of 7SK DMS reactivity profiles from bulk probing of SRSF1-APEX2 and LMNA-APEX2 cell lines and previous published[60] 7SK reactivity profiles. Modest differences in overall reactivity magnitude can be explained by the different probing conditions. In the current study, HEK293T cell lines expressing APEX fusions were probed using 2% DMS for 3 min at room temperature. In our prior study, HEK293T cells were probed using 1.6% DMS for 6 min at 37°C.
- **C.** Heatmap of pairwise Pearson correlations between speckle and lamina datasets across biological replicates.
- **D.** DANCE deconvolution of bulk 7SK under RAID-MaP conditions resolves expected A, B, H structural equilibrium. States are assigned as A, B, and H based on their deconvoluted reactivity profiles and correspondence to previously characterized 7SK conformations[54,60]. Colored arcs represent base-pairing probabilities derived from co-mutation rates, with green indicating the highest probability, followed by yellow and blue, and gray indicating the lowest probability.
- **E.** ΔR profiles of speckle- and lamina-enriched 7SK. Bar plots show mean ΔR ± s.e.m. for speckle-enriched ($n = 3$) and lamina-enriched ($n = 2$) samples.
- **F.** Windowed mean ΔR for 7SK RNA in lamina. Nucleotides more reactive (red) or less reactive (blue) in lamina-enriched samples are plotted on a symlog scale. Black outlined circles denote nucleotides with statistically significant differences ($p < 0.05$, Wilcoxon rank-sum test).

**Table S1: Oligos and primers**

| | |
|---|---|
| **gBlock for *in vitro* transcription (IVT)** | |
| AdenosylCobalamin riboswitch | TTCTAATACGACTCACTATAGGCCAAAACAACAACCGGTTAAAGCCTTATGGTCGCTACCATTGCACTCCGGTAGCGTTAAAAGGGAAGACGGGTGAGAATCCCGCGCAGCCCCCGCTACTGTGAGGGAGGACGAAGCCCTAGTAAGCCACTGCCGAAAGGTGGGAAGGCAGGGTGGAGGATGAGTCCCGAGCCAGGAGACCTGCCATAAGGTTTTAGAAGTTCGCCTTCGGGGGGAAGGTGAACAAAAACCAAACCAAAGAAACAACC |
| **Reverse-transcription and sequencing primers** | |
| AdenosylCobalamin riboswitch - RT | CCTACACGACGCTCTTCCGATCTTNNNNNNNNNNNGGTTGTTTCTTTGGTTTGGTTTTG |
| AdenosylCobalamin riboswitch – PCR1-F | GACTGGAGTTCAGACGTGTGCTCTTCCGATCTNNNNNGGCCAAAACAACAACCGG |
| AdenosylCobalmin riboswitch – PCR1-R | CCTACACGACGCTCTTCCGATCTT |
| 7SK-RT | CCTACACGACGCTCTTCCGATCTTNNNNNNNNNNNAAGAAAGGCAGAC |
| 7SK-PCR1-F | GACTGGAGTTCAGACGTGTGCTCTTCCGATCTNNNNNGGATGTGAGGGCGATCTGG |
| 7SK-PCR1-R | CCCTACACGACGCTCTTCCGATCT |
| **qPCR primers** | |
| MTCO2-qPCR-F | AACCAAACCACTTTCACCGC |
| MTCO2-qPCR-R | CGATGGGCATGAAACTGTGG |
| GAPDH-qPCR-F | GTCAACGGATTTGGTCGTATTG |
| GAPDH-qPCR-R | TGTAGTTGAGGTCAATGAAGGG |
| MALAT1-qPCR-F | GACGGAGGTTGAGATGAAGC |
| MALAT1-qPCR-R | ATTCGGGGCTCTGTAGTCCT |
| **ASOs * = phosphorothioate, 2MOE = 2'-O-methoxy-ethyl, /5Hexynyl/ = 5' Hexynyl** | |
| Scramble | /5Hexynyl/*/i2MOErA/*/i2MOErA/*/i2MOErT/*/i2MOErG/*/i2MOErG/*/i2MOErA/*/i2MOErA/*/i2MOErT/*/i2MOErG/*/i2MOErG/*/i2MOErA/*/i2MOErA/*/i2MOErT/*/i2MOErG/*/i2MOErG/*/i2MOErA/*/i2MOErA/*/i2MOErT/*/i2MOErG/*/32MOErG/ |
| Site-A0 | /5Hexynyl/*/i2MOErG/*/i2MOErA/*/i2MOErT/*/i2MOErC/*/i2MOErC/*/i2MOErT/*/i2MOErC/*/i2MOErC/*/i2MOErC/*/i2MOErC/*/i2MOErC/*/i2MOErG/*/i2MOErA/*/i2MOErC/*/i2MOErT/*/i2MOErC/*/i2MOErG/*/i2MOErG/*/i2MOErA/*/32MOErA/ |
| Domain 0/Core | /5Hexynyl/*/i2MOErA/*/i2MOErA/*/i2MOErG/*/i2MOErA/*/i2MOErC/*/i2MOErG/*/i2MOErG/*/i2MOErG/*/i2MOErT/*/i2MOErC/*/i2MOErG/*/i2MOErG/*/i2MOErG/*/i2MOErT/*/i2MOErG/*/i2MOErG/*/i2MOErG/*/i2MOErT/*/i2MOErA/*/i2MOErG/*/32MOErC/ |
| Domain IV | /5Hexynyl/*/i2MOErC/*/i2MOErC/*/i2MOErT/*/i2MOErG/*/i2MOErG/*/i2MOErT/*/i2MOErC/*/i2MOErC/*/i2MOErG/*/i2MOErC/*/i2MOErA/*/i2MOErC/*/i2MOErC/*/i2MOErA/*/i2MOErG/*/i2MOErT/*/i2MOErT/*/i2MOErC/*/i2MOErT/*/i2MOErA/*/32MOErA/ |
| **FISH probe** | |
| 7SK | GTGTCTGGAGTCTTGGAAGC/3Cy3Sp/ |
